# DAXX safeguards heterochromatin formation in embryonic stem cells

**DOI:** 10.1101/2021.04.28.441827

**Authors:** Antoine Canat, Adeline Veillet, Renaud Batrin, Clara Dubourg, Robert Illingworth, Emmanuelle Fabre, Pierre Therizols

## Abstract

Genomes comprise a large fraction of repetitive sequences folded into constitutive heterochromatin to protect genome integrity and cell identity. *De novo* formation of heterochromatin during preimplantation development is an essential step for preserving the ground-state of pluripotency and the self-renewal capacity of embryonic stem cells (ESCs). Yet, the molecular mechanisms responsible for the remodeling of constitutive heterochromatin are largely unknown. Here, we find that DAXX, an H3.3 chaperone, is essential for ESCs maintenance in the ground-state of pluripotency. DAXX accumulates at pericentromeric regions, and recruits PML and SETDB1, thereby promoting heterochromatin formation. In absence of DAXX or PML, the 3D-architecture and physical properties of pericentric and peripheral heterochromatin are disrupted, resulting in derepression of major satellite DNA, transposable elements and genes associated with the nuclear lamina. Using epigenome editing tools, we observe that H3.3, and specifically H3.3K9 modification, directly contribute to maintaining pericentromeric chromatin conformation. Altogether, our data reveal that DAXX is crucial for the maintenance and 3D-organization of the heterochromatin compartment and protects ESCs viability.

## Introduction

Eukaryotic nuclei generally contain two major genomic compartments, referred to as A and B, which respectively contain the active and inactive fractions of the genome. At the chromosome level, these compartments are characterized by alternating gene-rich and gene-poor domains. The B compartment is composed primarily of repetitive sequences, with satellite tandem repeats comprising one of the most abundant classes. Pericentromeric regions in mouse are composed of tandem repeats of a 234 bp sequence, named Major Satellite, that constitute approximately 8 Mb on each chromosome (Consortium et al., 2002; Guenatri et al., 2004). Most genes and repeated sequences of the B compartment are transcriptionally repressed by cooperative epigenetic mechanisms leading to the formation of constitutive heterochromatin. The three-dimensional organization of constitutive heterochromatin is an important layer regulating transcription (Falk et al., 2019; Lu et al., 2021). In some species, including the mouse, the pericentromeric heterochromatin (PCH) of different chromosomes aggregates to form large DAPI-dense heterochromatin clusters, called chromocenters. Disruption of satellite clustering is often associated with increased DNA damage and defects in chromosome segregation that are seen in pathologies such as Alzheimer’s disease and breast cancer (Hahn et al., 2013; Jagannathan et al., 2018; Mansuroglu et al., 2016; Zhu et al., 2011). Gene-poor domains are preferentially found beneath the nuclear edge and often referred as Lamina-Associated Domains (LADs) (Guelen et al., 2008). The tethering to the nuclear envelope is stochastic, and LADs are also found associated with other nuclear landmarks such as chromocenters and nucleolus (Koningsbruggen et al., 2010; Wijchers et al., 2015).

At the molecular level, constitutive heterochromatin exhibits covalent modifications including DNA methylation, H3K9 and H4K20 trimethylation (H3K9me3 and H4K20me3). Maintenance of H3K9me3 at pericentromeric satellites relies on the SUV39H1/2 methyltransferases, whereas a distinct H3K9me3 methyltransferase, SETDB1, operates at dispersed DNA repeats and telomeres (Fukuda et al., 2018; Gauchier et al., 2019; Martens et al., 2005; Matsui et al., 2010).

Heterochromatin maintenance is intrinsically linked to chromatin replication during S phase, but also requires the incorporation of histone variants, such as macroH2A and H3.3, into chromatin independently of DNA synthesis (Buschbeck and Hake, 2017; Mendiratta et al., 2019). The histone variant H3.3 differs from the canonical histone H3 by 4 or 5 residues and is deposited throughout the cell cycle in active and inactive chromatin by different histone chaperones. These histone-binding proteins prevent promiscuous incorporation into chromatin (Szenker et al., 2011). Together with ATRX, DAXX forms a histone chaperone complex responsible for H3.3 incorporation into heterochromatin regions, including PCH, retrotransposons and telomeres (Drané et al., 2010; Elsässer et al., 2015; He et al., 2015; Lewis et al., 2010; Sadic et al., 2015). At telomeres and retrotransposons, DAXX promotes heterochromatin formation by recruiting SUV39H1/2 or SETDB1, hereby facilitating the deposition of H3K9me3 (Elsässer et al., 2015; Gauchier et al., 2019; He et al., 2015; Hoelper et al., 2017). In addition to its role in H3.3 deposition, DAXX also prevents promiscuous incorporation of H3.3 by recruiting soluble H3.3-H4 dimers to PML nuclear bodies, membrane-less hubs for post-translational modification of proteins (Delbarre et al., 2013; Delbarre et al., 2017; Lallemand-Breitenbach and Thé, 2018).

Heterochromatin is *de novo* established during early embryogenesis, involving an important remodeling of its molecular composition and 3D-organization (Burton and Torres-Padilla, 2014). However, the underlying mechanisms and their functional relevance for early development remain elusive. During preimplantation development, the DNA methylation inherited from parental gametes is erased and reaches its minimal level in the blastocyst, defining the ground-state of pluripotency (Leitch et al., 2013; Ying et al., 2008). This wave of DNA demethylation directly leads to the transcriptional upregulation of several repetitive sequences, a process that is essential for the remodeling of constitutive heterochromatin (Jachowicz et al., 2017; Lu et al., 2021; Probst et al., 2010). For instance, the upregulation of major satellite transcripts is required for PCH reorganization into chromocenters and normal developmental progression (Casanova et al., 2013; Probst et al., 2010). Yet, derepression of DNA repeats correlates with high DNA damage signaling and their prolonged expression results in developmental arrest, suggesting that silencing through a DNA methylation-independent mechanism is required for normal embryogenesis (Jachowicz et al., 2017; Ziegler-Birling et al., 2009). Polycomb group proteins were previously shown to bind and facilitate silencing of repetitive sequences upon DNA hypomethylation (Saksouk et al., 2014; Tosolini et al., 2018; Walter et al., 2016). However, knockout of most polycomb-related genes does not impact pre-implantation embryogenesis (Aloia et al., 2013), implying that other factors maintain heterochromatin in the absence of DNA methylation.

In mouse, both DAXX and H3.3 deletions cause genomic instability resulting in early embryonic lethality with most knockout embryos failing to reach the blastocyst stage (Jang et al., 2015; Liu et al., 2020; Michaelson et al., 1999). DAXX binds major satellite regions during the earliest stages of development, just before PCH reorganizes to form chromocenters (Arakawa et al., 2015; Liu et al., 2020). While the importance of DAXX for chromocenter formation remains unknown, the viability of *Daxx^-/-^* embryos drops after chromocenter formation (Liu et al., 2020; Probst et al., 2010), suggesting a direct link between DAXX function at chromocenters and developmental progression. Surprisingly, *Daxx* is not required for *in vitro* culture of blastocyst-derived Embryonic Stem Cells (ESCs) (Elsässer et al., 2015; He et al., 2015; Hoelper et al., 2017). In ESCs, DAXX is not recruited to PCH, but rather to retrotransposons and telomeres (Elsässer et al., 2015; Gauchier et al., 2019; He et al., 2015; Saksouk et al., 2014). However, the recruitment of DAXX to chromatin is tightly associated with the low level of DNA methylation that characterize preimplantation embryogenesis (Arakawa et al., 2015; He et al., 2015; Liu et al., 2020). ESCs are typically cultured in serum-based media and accumulate high levels of DNA methylation (Leitch et al., 2013), which could explain the discrepancies with *in vivo* observations. ESCs can maintain low levels of DNA methylation and reach the ground-state of pluripotency when grown in a serum-free medium, named 2i, but the role of DAXX for their survival and the formation of PCH has yet to be addressed.

Here, we decipher the role played by DAXX at PCH in pluripotent ESCs. We find that DAXX is essential for ground-state ESC survival. We provide evidence that DAXX localizes to pericentric heterochromatin and recruits PML and SETDB1, facilitating heterochromatin formation and organization through H3.3K9 modification. The deletion of *Daxx* or *Pml* impacts pericentromeric heterochromatin formation altering its biophysical and clustering properties. Thus, our results identify DAXX as an essential regulator preserving heterochromatin integrity and maintaining the viability of ground-state ESCs.

## Results

### DAXX is essential for ESC survival upon ground-state conversion

To assess the role of DAXX in pluripotent cells, we generated a *Daxx* knock-out (KO) ESC line. We targeted exon 3 of *Daxx* using CRISPR/cas9 technology. While we confirmed the absence of DAXX mRNA and protein (Fig. 1A; Fig. S1A), there were no obvious changes in H3.3 protein levels in the resulting *Daxx* KO ESC line (Fig. 1A). Consistent with previous reports, loss of DAXX did not impact the growth of ESCs cultured with a serum-based medium (Elsässer et al., 2015). Likewise, neural differentiation induced by leukemia inhibitory factor withdrawal and retinoic acid addition did not impact on cell viability in *Daxx* KO ESC (Diff., Fig. 1B). When the *Daxx* KO and WT ESCs were converted to the ground state, using a medium with two small kinase inhibitors and vitamin C (hereafter denoted 2iV), no growth defect was detected for *Daxx* KO cells after 4 days of conversion, but prolonged culture in 2iV medium induced a drastic decrease in cell viability (Fig. 1B). Compared to the parental WT ESCs, only 12.6% of *Daxx* KO ESCs remained after 8 days of culture in 2iV (Fig. 1B). No surviving cells could be detected after 9 to 10 days of conversion. We confirmed that under serum conditions, ESCs have higher levels of DNA methylation than the blastocyst cells from which they are derived (Fig. S1B) (Borgel et al., 2010). In contrast, ESCs cultured in 2iV, exhibit very low levels of DNA methylation (Blaschke et al., 2013). Altogether, these results indicate an essential role for DAXX in the maintenance of pluripotent cell survival upon ground-state conversion.

**Fig. 1.**
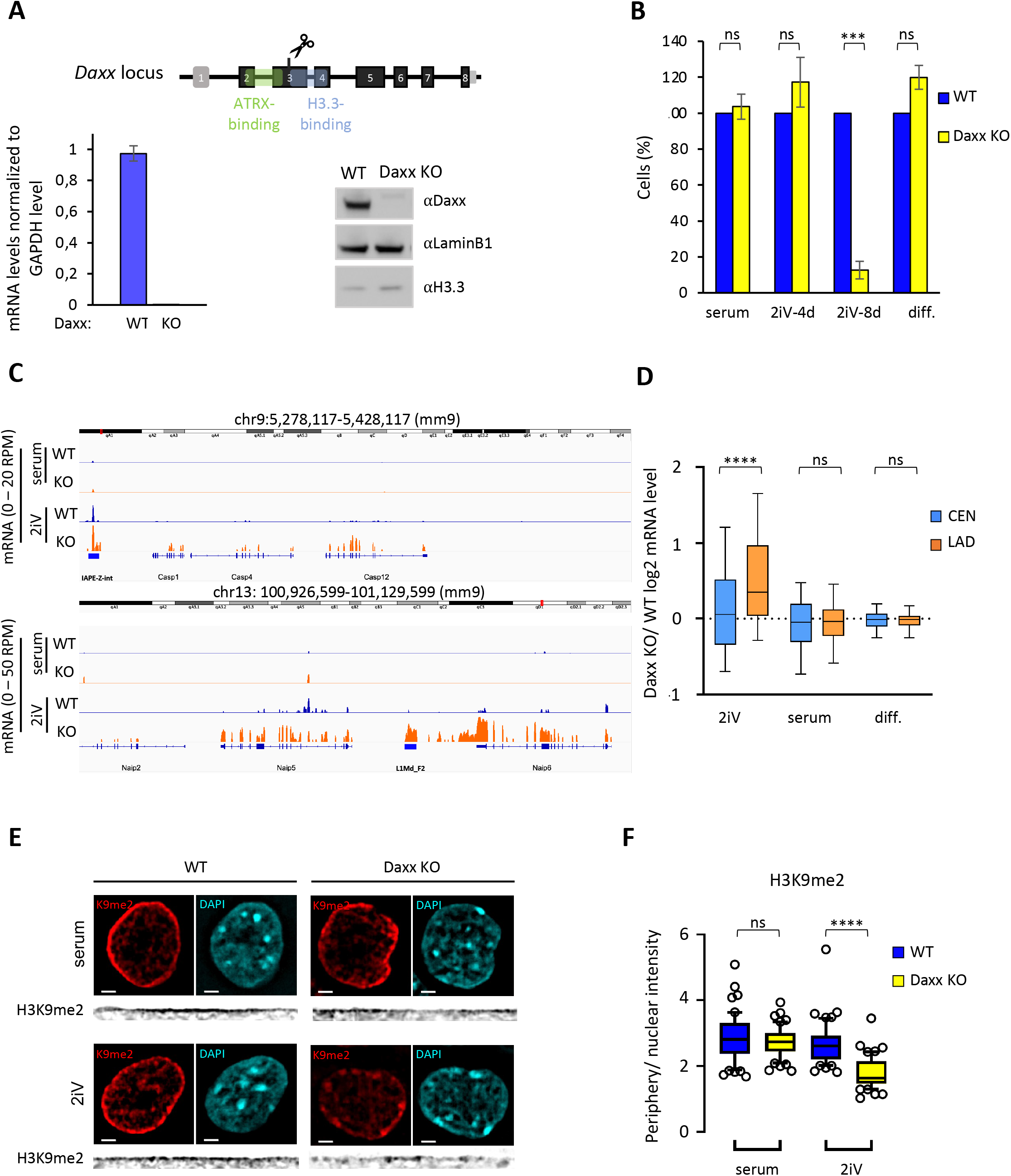
DAXX is essential for ESC survival upon ground-state conversion and is required for transcriptional repression of peripheral heterochromatin. **(A)** Schematic of the *Daxx* locus and the of the guide RNA targeting exon 3. Left, RT-qPCR experiment on DAXX RNA in WT cells or cells after CrispR/cas9 editing on *Daxx.* The mean of two biological replicates with SEM are shown. Right, western blot with anti-DAXX antibody, anti-LAMIN-B1 and anti-H3.3 served as loading control. One experiment from at least 2 biological replicates is shown. **(B)** Quantification of the number of cells (in %) compared to WT in each culture condition, serum, after 4 days (4d) or 8 days (8d) of 2iV conversion and after 5 days of Retinoic Acid differentiation (Diff.) Histogram represents mean with SEM of 3 biological replicates for each condition except Diff. (mean of two biological replicates). T-tests were used for statistical analysis. **(C)** RNA-seq tracks of 2 Lamin Associated Domains (LAD) of WT ESCs (blue) and *Daxx* KO (orange) under serum and 2iV conditions. **(D)** Boxplots of the log2 fold change of mRNA levels of genes located in LAD (LAD, orange) and non-LAD (CEN, blue) regions in *<u>Daxx</u>* KO over WT ESCs, grown in 2iV, serum and diff. media. Two-sided Mann-Whitney tests was used for statistical analysis **(E)** Representative nuclei after immuno-detection of H3K9me2 (red) and counterstaining with DAPI (blue) in WT and *Daxx* KO ESCs in serum and 2iV (scale bar = 2μm). Linescans below represent straighten H3K9me2 signal at the nuclear periphery (black). **(E)** Boxplots of H3K9me2 signal quantification enrichment at the nuclear periphery from 2 independent replicates. Two-sided Mann-Whitney tests was used for statistical analysis.

To further investigate the impact of *Daxx* knockout, we monitored the transcriptional landscapes under different conditions using RNA sequencing (RNA-seq). *Daxx* KO and WT cells show only negligible differential expression of protein coding genes upon differentiation with retinoic acid (Fig. S1C; Fig. S1D). We concluded that the absence of DAXX only has a minor impact on transcriptome changes upon neural differentiation. This result is consistent with the absence of morphological or cell growth defects observed upon differentiation and confirms that DAXX is dispensable for neuronal differentiation. After 4 days of 2iV conversion a substantial differential expression of genes in *Daxx* KO and WT ESCs was observed (Fig. S1C). However, the global transcriptional response of *Daxx* KO ESCs to 2iV medium was similar to WT ESCs (Fig. S1D). We compared the expression of 166 markers previously identified as differentially expressed upon 2iV conversion (Blaschke et al., 2013). For both cell lines most of these markers changed expression upon conversion according to the expected pattern (Fig. S1E) supporting the conclusion that *Daxx* KO ESCs can reach the ground-state of pluripotency after 4 days of 2iV conversion.

Overall, our data show that *Daxx* is essential for the survival of ground-state ESCs but their loss of viability is unlikely caused by a conversion defect. Despite the mis-regulation of over 2500 genes after 4 days of 2iV conversion, no significant gene network that could explain the subsequent loss of viability in *Daxx* KO ESCs could be identified.

### DAXX contributes to the transcriptional repression of DNA repeats and peripheral heterochromatin

As DAXX is known to be responsible for H3.3 deposition in heterochromatic regions, we investigated the impact of *Daxx* deletion on the transcriptional repression of the silenced portion of the genome (Elsässer et al., 2015).

LADs make up a significant portion of the silenced compartment. To investigate the effect of *Daxx* deletion on the expression of genes located in the 1180 LADs described in ESCs (Peric-Hupkes et al., 2010), we compared their differential expression to that of genes not interacting with the lamina (CEN). In serum and differentiated conditions, we found that LAD genes had very similar differential expression to genes not located in LADs (Fig. 1C; Fig. 1D). However, we observed an increase in the mRNA levels of several LAD genes normally silenced such as *Casp1, Casp4,* and *Casp12* on chromosome 9 and *Naip6* and *Naip5* on chromosome 13, in *Daxx* KO ESCs under 2iV condition (Fig. 1C; Fig. 1D; Fig. S2A). Genome-wide, the differential expression of genes located in LADs was significantly increased in the 2iV condition, suggesting that the presence of DAXX is necessary for proper silencing of these genes in ground-state ESCs.

Histone modification H3K9me2 is a hallmark of LADs and generally accumulates beneath the nuclear envelope (Kind et al., 2013; Peric-Hupkes et al., 2010). To assess whether the global upregulation of LAD genes observed in *Daxx* KO ESCs upon 2iV conversion was associated with a defect in peripheral heterochromatin assembly, we examined the accumulation of H3K9me2 by immunofluorescence. In WT and *Daxx* KO ESCs grown in serum condition, the H3K9me2 signal was irregularly diffuse in the nucleoplasm with a significant accumulation at the nuclear edge (Fig. 1E). Quantification of H3K9me2 distribution by linescan density did not show any significant difference in the enrichment of signal at the nuclear periphery between WT and *Daxx* KO ESCs in serum (Fig. 1F; Fig. S2B). However, after 4 days of 2iV conversion, the H3K9me2 signal remained enriched at the nuclear rim in WT converted cells, but only small patches along the nuclear periphery were observed in the absence of DAXX (Fig. 1E). Linescan quantification confirmed a significant decrease in H3K9me2 enrichment at the nuclear periphery in *Daxx* KO ESCs (Fig. 1F). In contrast, LaminB1 accumulated normally at the nuclear edge in both cell lines regardless of growing conditions, indicating no global nuclear lamina assembly defects in *Daxx* KO ESCs (Fig. S2C; Fig. S2D). We therefore conclude that DAXX is necessary for the maintenance of peripheral heterochromatin.

Overall, our observations show that *Daxx* deletion impacts the regulation of the nuclear periphery by altering the tethering of peripheral heterochromatin and the transcriptional silencing of genes located in LADs.

### DAXX relocates to chromocenters upon ground-state conversion

We additionally found that many endogenous retrovirus families were upregulated in *Daxx* KO ESCs in serum and 2iV (Fig. 2A; Fig. S3), consistent with previous observations (Elsässer et al., 2015; He et al., 2015). Several LINE1 elements, notably the L1MdT and L1MdA families, not previously reported as DAXX targets, were also upregulated in pluripotent *Daxx* KO ESCs (Fig. 2A; Fig. S3). The role of DAXX on transcriptional silencing was not limited to interspersed repeats as we detected a strong upregulation of major satellites RNA in both serum and 2iV condition in the *Daxx* KO cells (Fig. 2A; Fig.S3).

**Fig. 2.**
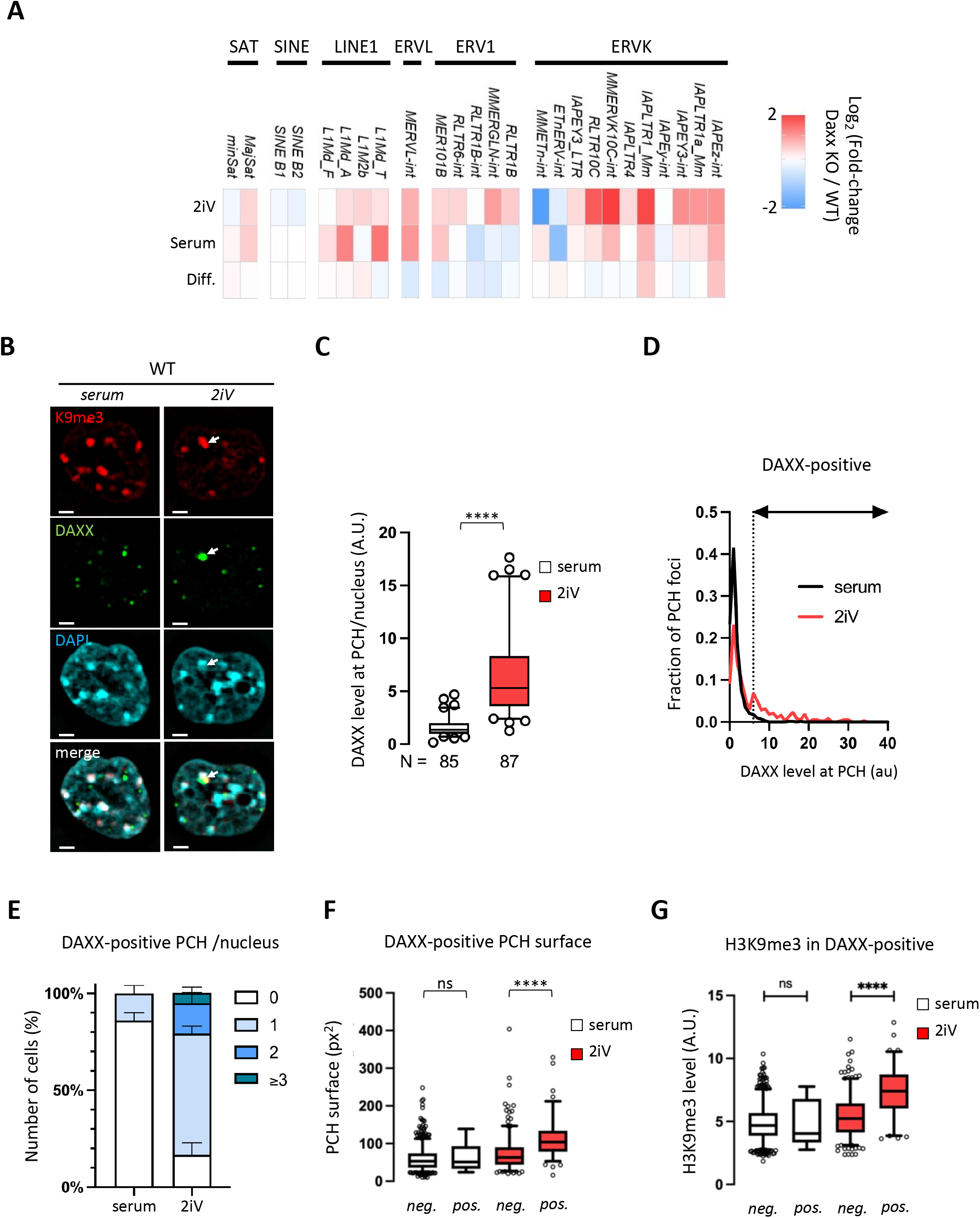
DAXX is recruited to PCH in 2iV converted ESCs. **(A)** Heatmap for different classes of transposable elements. Data are mean from RNA-seq experiments on 3 biological replicates. **(B)** Representative nuclei of ESCs grown in serum or 2iV immuno-detected with H3K9me3 (red) and DAXX (green). Nuclei were counterstained with DAPI (cyan). Scale bar = 2μm. **(C)** Boxplot of mean DAXX levels at pericentromeric heterochromatin (PCH) in WT ESCs grown in serum (white) or in 2iV (red) from 3 biological replicates. **(D)** Distribution curve of DAXX intensity observed at individual PCH foci in WT ESCs grown in serum (white) or in 2iV (red) from 3 independent replicates. Dashed line indicates the threshold used to defined DAXX-positive PCH. **(E)** Histogram of the average fraction of the cell population with 0, 1, 2 and 3 or more, DAXX-positive PCH foci in WT ESCs in serum and 2iV. SEM are shown. **(F)** Boxplots of DAXX-negative (neg) and DAXX-positive (pos) PCH foci surface in WT ESCs grown in serum (white) or in 2iV (red). **(G)** Boxplots of H3K9me3 intensity in DAXX-negative (neg) and DAXX-positive (pos) PCH foci in WT ESCs grown in serum (white) or in 2iV (red). Mann-Whitney tests were used for statistical analysis in C, F and G.

To investigate whether DAXX binds to major satellites in ESCs, we used immunofluorescence with H3K9me3 staining to track PCH after confirming the strong correlation between H3K9me3 and major satellites using immuno-FISH (Fig. S4A; Fig. S4B). In both serum and 2iV conditions, DAXX signal concentrated in the nucleoplasm as round foci (Fig. 2B; Fig. S4C). The distribution of foci sizes was similar in both growing conditions, but a small fraction of larger foci could be observed under 2iV condition. We therefore set a threshold defining small and larger DAXX foci (Fig. S4D). The large DAXX foci exhibited higher levels of DAPI and H3K9me3 specifically under 2iV condition, supporting that DAXX can relocate to PCH in ground-state ESCs (Fig. S4E; Fig. S4F). To confirm this observation, we segmented nuclei and PCH foci using DAPI and H3K9me3 signals, respectively, allowing us to measure the enrichment of DAXX signal at PCH per nuclei (Fig. S4G). The level of DAXX observed at PCH upon 2iV conversion was significantly higher than in the serum condition (Fig. 2C). Notably, the increase in DAXX enrichment in 2iV was not homogenous among the H3K9me3 foci population, but rather the result of a subset of clusters showing strong DAXX enrichment, allowing us to determine a threshold to define DAXX-positive PCH (Fig. 2D). While only 14% of cells showed one DAXX-positive PCH cluster in the serum condition, 83% of 2iV-converted cells had at least one DAXX-positive H3K9me3 focus (Fig. 2E). A small proportion of cells showed 2 or 3 or more DAXX-enriched PCH foci (16% and 5%, respectively). Interestingly, our analysis revealed that DAXX-enriched PCH foci were bigger in size and had higher H3K9me3 staining intensities, suggesting a potential link between DAXX and H3K9me3 regulation (Fig. 2F; Fig. 2G).

Altogether, our data show that DAXX accumulates in a subset of PCH clusters in ground-state ESCs, suggesting that it may play a role in the maintenance of PCH and the transcriptional silencing of major satellite sequences.

### DAXX maintains pericentric heterochromatin organization in pluripotent cells

To investigate the role that DAXX might play at PCH in ground-state ESCs, we next investigated whether DAXX facilitates PCH formation in ESCs. Since the loss of pericentromeric silencing is generally caused by defective heterochromatin assembly and often correlates with impaired clustering of chromocenters (Hahn et al., 2013; Healton et al., 2020; Pinheiro et al., 2012; Zhu et al., 2011), we performed DNA FISH experiments against the major satellites and segmented the 3D signal to determine the number of clusters (Fig. 3A). In serum or differentiated cells, the deletion of *Daxx* had no significant impact on the number of clusters (Fig. 3B). However, after 4 days of 2iV conversion, the average number of major satellite foci was increased by 37% in the *Daxx* KO ESCs. The presence of an increased number of PCH foci indicates that DAXX is required for their clustering in ESCs, specifically under 2iV condition and correlates with the loss of viability observed in these cells at later stages of conversion (Fig. 1B). To confirm the role of DAXX in PCH clustering, we used the TALE (transcription activator–like effector) epigenome editing tool to ask whether direct DAXX recruitment to major satellites was sufficient to reduce the number of foci. We generated a FLAG-tagged TALE engineered to specifically target major satellite sequences (hereafter called TALE_MajSat_), following the design of a previous study (Miyanari et al., 2013). Immunofluorescence against the FLAG tag of the TALE_MajSat_ confirmed its colocalization with the major satellite FISH signal (Fig. 3C). When bound to pericentromeres in pluripotent *Daxx* KO cells, TALE_MajSat_-DAXX restored chromocenter clustering with the formation of fewer PCH foci (Fig. 3D). In WT ESCs, artificial tethering of DAXX to chromocenters had a similar impact, increasing endogenous PCH clustering (Fig. S5A). These results indicate that DAXX enhances physical interactions between major satellite sequences and contributes to the 3D organization of PCH.

**Fig. 3.**
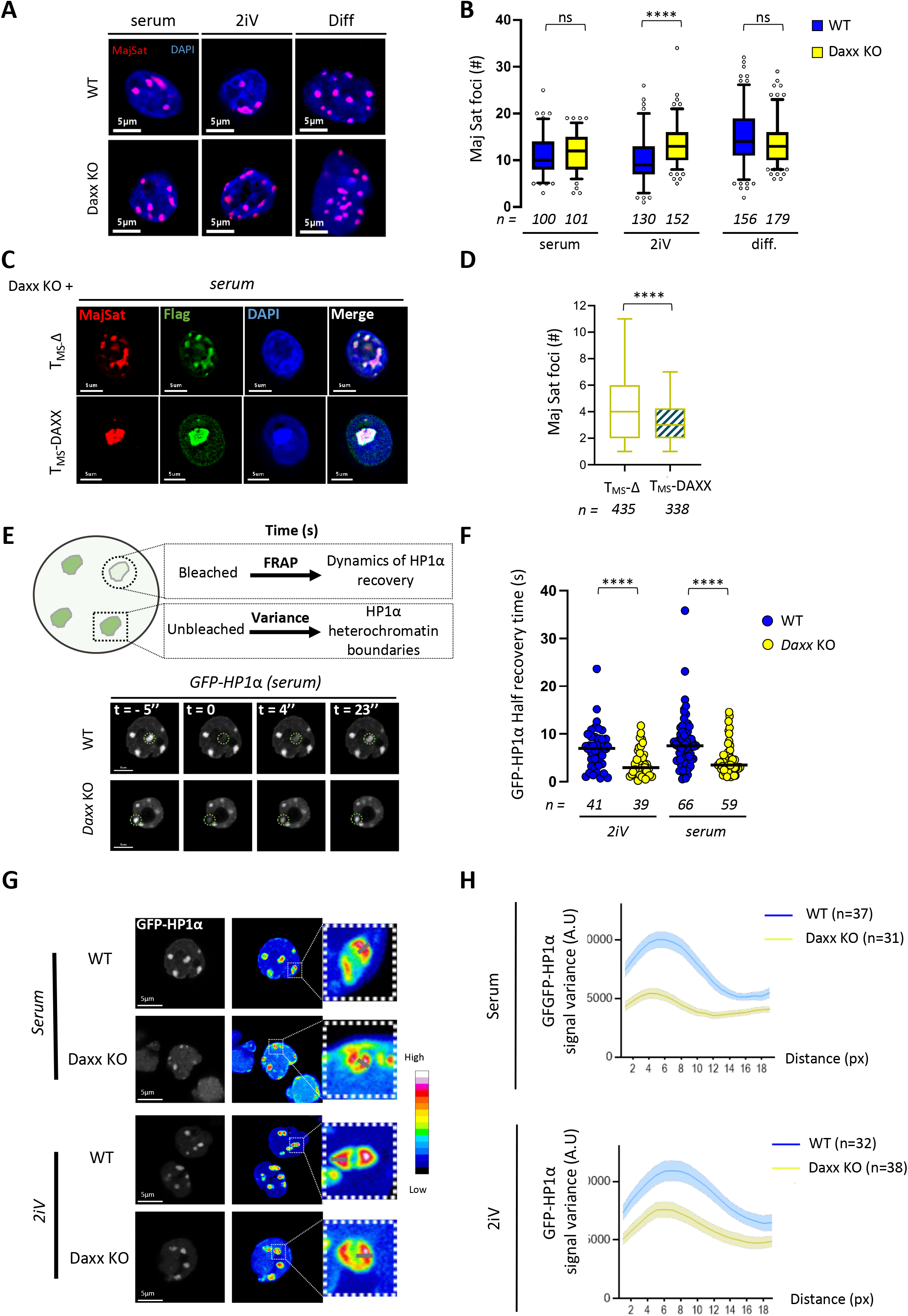
DAXX maintains heterochromatin organization in pluripotent cells. **(A)** Representative major satellite (MajSat) DNA FISH pictures of WT and Daxx KO ESCs in serum, 2iV and differentiation (Diff.) conditions. **(B)** Quantification of the number of major satellite foci per nucleus in WT (blue) or *Daxx* KO (yellow) ESC. n=total number of nuclei analyzed from at least 3 biological replicates. **(C)** Representative immunoFISH pictures of major satellite and Flag for *Daxx* KO cells transfected with TALE_MajSat_-Δ or TALE_MajSat_-DAXX. **(D)** Quantification of the number of major satellite foci detected in the focal plane. n= total number of nuclei analyzed from 4 biological replicates. **(E)** Top, scheme depicting the two different analyses used in live-imaging experiments. Bottom, representative pictures from FRAP experiments performed on WT and *Daxx* KO cells transfected with GFP-HP1α. t=-5’’ corresponds to pre-bleach fluorescence. t=0 corresponds to the laser bleach pulse. t=4’’ and t=23’’ correspond to post-bleach recovery images after 4 and 23 seconds respectively. **(F)** Quantification of half-recovery times in seconds for individual nuclei. n=total number of nuclei analyzed from 2 biological replicates. **(G)** Representative pictures of GFP-HP1α and its corresponding variance in fluorescence intensities along time in serum or 2iV, WT or *Daxx* KO ESCs. White dashed squares display magnification of individual chromocenters. Color correspondence of signal variance strength (from high to low) is shown. Scale bar, 5μm. **(H)** Graph displaying the variance intensities along measured on 1μm lines traced above non-bleached foci. Two-sided Mann-Whitney tests were used for statistical analysis in B, D and F.

We hypothesized that the decreased chromocenter clustering seen in *Daxx* KO ESCs might result from defective heterochromatin formation. Functional heterochromatin forms a self-segregating subcompartment with limited protein exchange with the rest of the nucleoplasm (Hinde et al., 2015; Strom et al., 2017). To test whether DAXX is important for heterochromatin segregation, we followed GFP-HP1α using live-imaging in WT and *Daxx* KO ESCs (Fig. 3E). We monitored the mobility of GFP-HP1α by measuring its fluorescence recovery rate at chromocenters after photobleaching (Fig. 3E; Fig. S5B). Compared to WT, the half-recovery time was significantly shorter in *Daxx* KO cells both in serum and 2iV conditions, suggesting higher protein exchange between chromocenter and the nucleoplasm in the absence of DAXX (Fig. 3F). However, the mobile fraction of GFP-HP1α remained constant in both cell lines, suggesting that the same amount of protein was bound to chromatin (Fig. S5B).

The boundary property of PCH is known to cause high variance in HP1α signal over time at the edge of a chromocenter (Strom et al., 2017). We reasoned that this higher HP1α recovery rate might arise from altered heterochromatin acting as a barrier to protein diffusion. We thus quantified temporal changes in GFP-HP1α signal intensity variation at unbleached chromocenters (Fig. 3G). We found that variance levels were low in the nucleoplasm of WT ESCs and only increased at chromocenters, peaking at their borders (Fig. 3G; Fig. 3H), confirming previous observations (Strom et al., 2017). However, in *Daxx* KO cells, the pattern of the variance from HP1α signal was altered. In both serum and 2iV conditions, the peak of variance at the edge of chromocenters was significantly lower in the absence of DAXX, suggesting a compromised heterochromatin barrier (Fig. 3H). Moreover, molecular analysis using MNase digestion revealed that *Daxx* deletion in pluripotent ESCs increased the accessibility of pericentric chromatin, indicating that PCH exists in an abnormal state in these cells (Fig. S5C).

Altogether, our data suggests that DAXX plays a crucial role in regulating PCH compaction state, chromocenter boundary properties, and major satellite clustering. In conclusion, DAXX is essential for proper assembly and spatial organization of pericentromeric heterochromatin in pluripotent ESCs.

### PML and DAXX share similar roles in ESCs

PML, identified as a partner of DAXX, has been recently shown to contribute to the transcriptional repression of transposable elements in ESCs (Tessier et al., 2022). To investigate the role of PML in the organization of the heterochromatic compartment in ground-state ESCs, we first analyzed its localization in relation to DAXX and H3K9me3 by immunofluorescence (Fig. 4A; Fig. S6A). Independently of culture conditions, PML formed small round foci within the nucleoplasm that colocalized with DAXX, known as nuclear bodies (Fig. 4A; Fig. S6A). Upon 2iV conversion, a fraction of larger PML foci were observed with higher levels of H3K9me3 and DAPI (Fig. S6B; Fig. S6C; Fig. S6D). These larger PML foci accumulated around PCH in the form of either complete or partial structures, referred to as PML-rim or PML-arc, respectively (Fig. 4A; Fig. S6E). Both the PML-rim and PML-arc structures were consistently associated with the presence of DAXX at PCH and were observed in similar proportions as DAXX-positive PCH clusters (Fig. 2E; Fig. 4B). This indicates that PML is recruited to PCH alongside DAXX in ground-state ESCs.

**Fig. 4.**
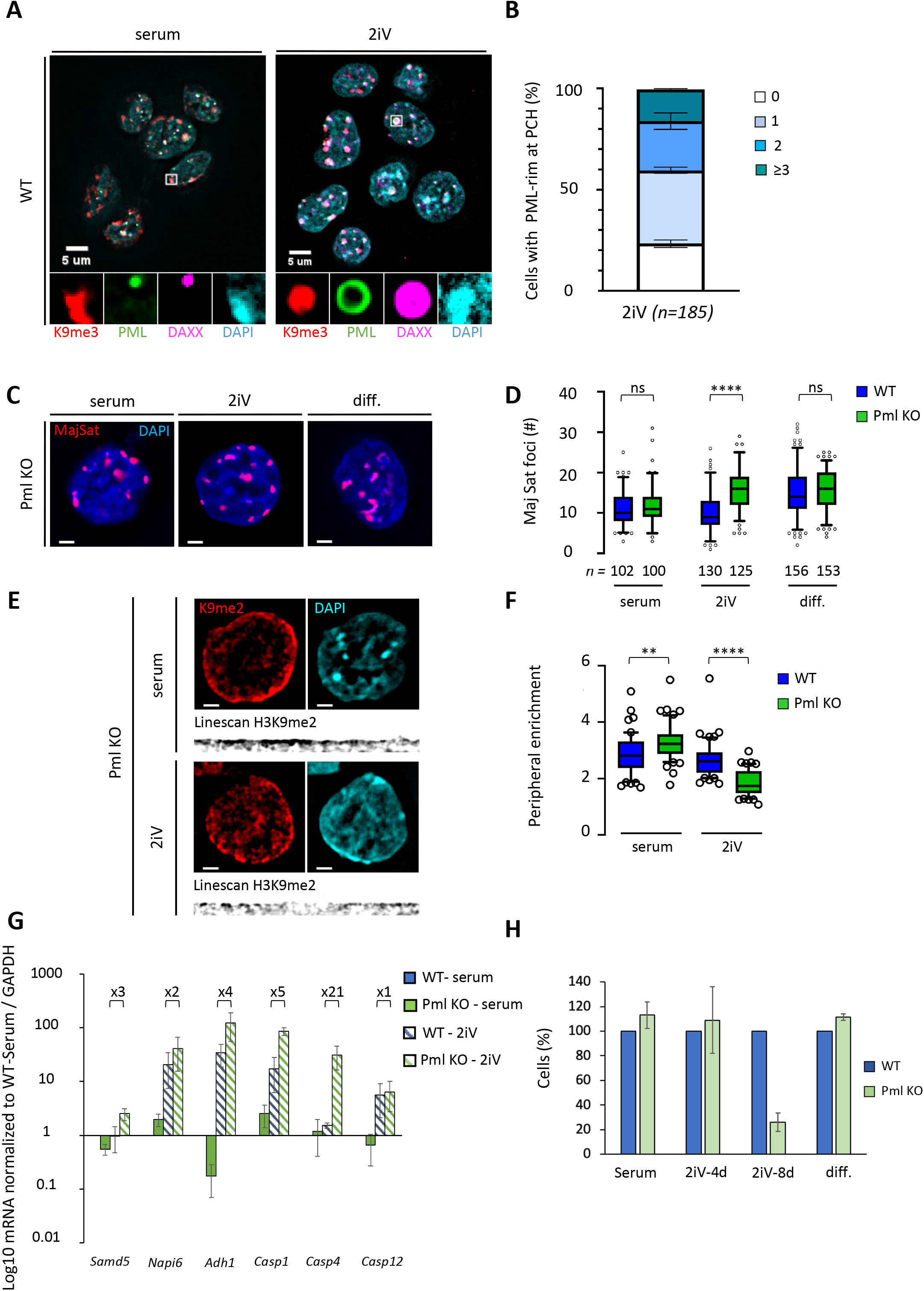
PML protects heterochromatin in ground-state ESCs. **(A)** Representative field of WT ESCs grown in serum or 2iV immuno-detection of H3K9me3 (red), PML (green), DAXX (purple) and counterstaining with DAPI (cyan). White squares highlight the absence (serum) or the presence (2iV) of PML-rim around H3K9me3 foci. **(B)** Histogram of the average fraction of cell population showing either 0, 1, 2 and 3 or more, PML-rims, with SEM, in 2i converted WT ESCs (n= total number of nuclei). **(C)** Representative major satellite DNA FISH pictures of *Pml* KO ESCs in serum, 2iV and differentiation (dif.) conditions. **(D)** Quantification of the number of major satellite foci per nucleus. WT values (blue) are taken from Fig. 3B. n=total number of nuclei analyzed from at least 3 biological replicates. **(E)** Representative nuclei immuno-detection of H3K9me2 (red) and counterstaining with DAPI (blue) in *Pml* KO EScs grown in serum and 2iV (scale bar = 2μm). Linescan represent straighten H3K9me3 signal at the nuclear periphery (black). **(F)** Boxplots of H3K9me2 signal quantification enrichment at the nuclear periphery. WT values (blue) are taken from Fig. 1F. n=total number of nuclei analyzed from at least 3 biological replicates. **(G)** RT-qPCR RNA quantification of LAD genes in WT and *Pml* KO ESCs in serum and upon 2iV conversion. Histogram represents means normalized to WT-serum with SEM of 3 biological replicates. T-tests were used for statistical analysis **(H)** Quantification of the number of cells compared to WT in each culture condition. Histogram represents mean with SEM of 3 biological replicates. Two-sided Mann-Whitney tests were used for statistical analysis in D and F.

To further examine the role played by PML at PCH clusters, we analyzed the distribution of major satellites in *Pml* KO ESC using FISH (Tessier et al., 2022). In *Pml* KO ESCs, no protein was detectable (Fig. S7A). Similar to the *Daxx* KO line, *Pml* KO ESCs showed a significant increase in the number of major satellite foci only after 4 days of 2iV conversion (Fig. 4C; Fig. 4D), suggesting that PML, like DAXX, is required for proper PCH clustering in ground-state ESCs.

The integrity of peripheral heterochromatin in *Pml* KO ESCs was analyzed. Immunofluorescence revealed that H3K9me2 accumulated at the nuclear periphery to a greater extent in serum conditions in *Pml* KO cells compared to WT cells (Fig. 4E; Fig. 4F). However, after 4 days of 2iV conversion, the H3K9me2 signal in *Pml* KO cells was similar to that observed in *Daxx* KO cells, with only small patches of H3K9me2 present beneath the nuclear envelope and a significant reduction in peripheral enrichment of the signal (Fig. 4F). In contrast, LaminB1 signal was not altered by Pml deletion supporting the conclusion that it is specific to peripheral heterochromatin (Fig. S7B; Fig. S7C). To determine the effect of this alteration of peripheral heterochromatin on gene expression, we analyzed the transcriptional repression of LAD genes that were upregulated after *Daxx* deletion. Out of the six selected genes, five showed higher mRNA levels in 2iV-converted *Pml* KO ESCs compared to WT cells (Fig. 4G). Although the transcriptional upregulation was less severe in *Pml* KO cells than in *Daxx* KO cells, our results suggest that PML, like DAXX, is involved in the transcriptional silencing of LAD genes.

The deletion of *Pml* and *Daxx* have very comparable consequences on the heterochromatin of ESCs after 4 days of 2iV conversion. We thus wondered how *Pml* deletion would impact the viability of ESCs under prolonged 2iV conversion (Fig. 4H). After 4 days of conversion, the viability of *Pml* KO was not impacted, but was decreased by nearly 75% after 8 days. Like the deletion of *Daxx,* this effect was specific to prolonged 2iV conversion as the absence of PML had no consequence on the viability under serum condition or after retinoic acid differentiation.

Overall, our data show that PML and DAXX play similar roles in ground-state ESCs. Like DAXX, PML is recruited to PCH and its absence impairs the proper organization of the heterochromatic compartment, significantly impacting cell viability.

### DAXX recruits SETDB1 to chromocenters in ground-state ESCs

We have shown that in ground-state ESCs, DAXX and PML are recruited to major satellites thereby facilitating PCH formation. To further investigate the mechanism underlying the function of DAXX and PML, we searched for DAXX- and PML-interacting factors that could be recruited specifically to chromocenters. Amongst the DAXX interacting partners, SETDB1, an H3K9me3 methyltransferase, colocalizes with DAXX at PML nuclear bodies and is important for repetitive DNA transcriptional silencing (Cho et al., 2011; Karimi et al., 2011). SETDB1 partially colocalized with PML in a DAXX-independent manner in serum conditions (Fig. S8A). Under 2iV treatment, SETDB1 relocalized to PCH with DAXX and PML in 70% of WT ESCs (Fig. 5A; Fig. 5B; Fig. S8B). By contrast, PML and SETDB1 were only associated with PCH in 5 and 17% of *Daxx* KO cells, respectively (Fig. 5B). We conclude that DAXX is critical for PML and SETDB1 recruitment to chromocenters. To confirm that DAXX was sufficient for SETDB1 recruitment to PCH, we expressed TALE_MajSat_, fused to DAXX in *Daxx* KO ESCs. TALE-mediated DAXX binding to major satellite repeats drastically increased the amount of SETDB1 signal observed at chromocenters (Fig. S8C; Fig. S8D).

**Fig. 5.**
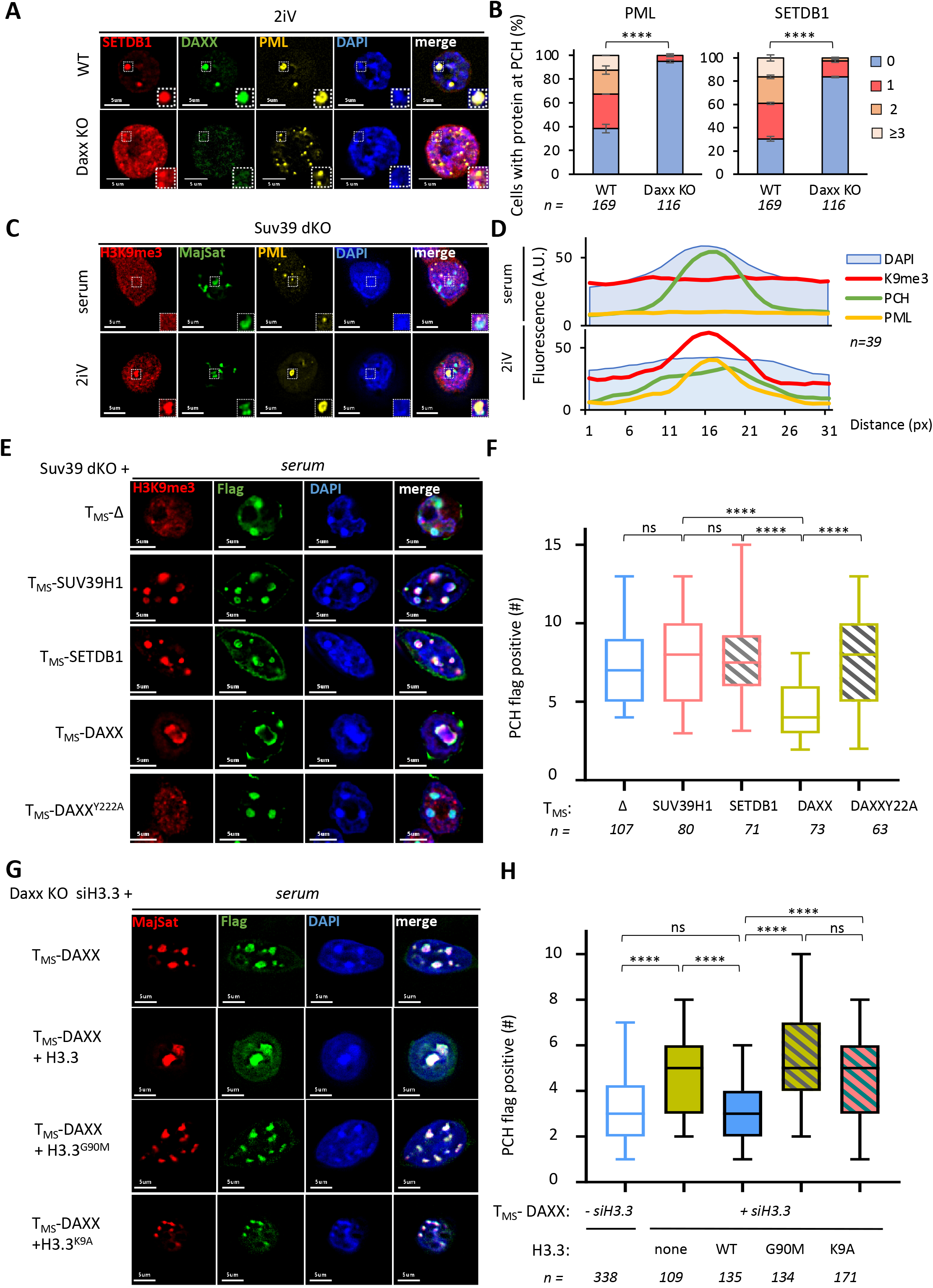
DAXX recruits Setdb1 and mediates chromocenter clustering via H3.3K9me3 modification. **(A)** Representative immunofluorescence pictures for SETDB1, DAXX and PML in WT or *Daxx* KO ESCs upon 2iV conversion. **(B)** Quantification of the mean number of PML or SETDB1 foci at chromocenters in WT and *Daxx* KO ESCs. n= total number of nuclei analyzed from two independent replicates. Chi-square tests were used for statistical analysis. **(C)** Representative immunoFISH pictures for H3K9me3, PML and major satellites in 2iV and serum of Suv39H1/2 dKO (double knock out) ESCs. White dashed squares highlight a major satellite focus. **(D)** Cumulated fluorescence intensities profile of H3K9me3 positive PCH foci in cells grown in serum or 2iV for either H3K9me3, PCH or PML signal. n=total number of nuclei analyzed from at least 2 biological replicates. **(E)** Representative immunofluorescence pictures for H3K9me3 and Flag in Suv39dKO serum ESCs transfected with TALE_MajSat_-Δ, TALE_MajSat_-SUV39H1, TALE_MajSat_-SETDB1, TALE_MajSat_-DAXX or TALE_MajSat_-DAXX^Y222A^. **(F)** quantification of the number of PCH foci observed in the different transfection conditions described in E. **(G)** Representative immunoFISH pictures for major satellite and Flag in *Daxx* KO serum ESCs co-transfected with siRNA against H3.3, TALE_MajSat_-DAXX and either no additional construct or H3.3WT, H3.3G90M or H3.3K9A. **(H)** Quantification of the number of major satellite foci detected of medium focal plane observed in the different transfection conditions described in G. For comparison, distribution T_MS_-DAXX without H3.3 knockdown data comes from Fig. 3D. n=total number of nuclei analyzed from 2 biological replicates. Two-sided Mann-Whitney tests were used for statistical analysis in F and H.

The presence of DAXX at PCH correlated with higher amounts of H3K9me3 in 2iV condition (Fig. 2G). However, DAXX deletion had no effect on H3K9me3 levels at chromocenters independently of culture conditions (Fig. S9A; Fig. S9B). Since DAXX deletion prevented SETDB1 recruitment to PCH, it suggests that the majority of H3K9me3 deposition at this locus relies on, or can be rescued by other lysine methyl transferases, such as SUV39H1/2 (Lehnertz et al., 2003; Peters et al., 2001).

To further investigate the conflicting results of DAXX-mediated SETDB1 recruitment and the unchanged amount of H3K9me3 in *Daxx* KO ESCs, we examined the individual roles of DAXX, SETDB1, and H3K9me3 in PCH spatial organization. We used Suv39H1/2 double knock-out (Suv39dKO) ESCs, in which H3K9me3 can only result from SETDB1 activity. Suv39dKO cells did not show any growth defects in 2iV medium, indicating the dispensability of both enzymes for pluripotency. Unlike WT ESCs, Suv39dKO did not accumulate H3K9me3 at major satellite foci, (Fig. S9A; Fig. 5C). However, we observed a strong accumulation of H3K9me3 in some major satellite foci surrounded by PML upon 2iV conversion, suggesting that the fraction of SETDB1 recruited to PCH in ground state is indeed functionally active (Fig. 5C; Fig. 5D). We next used the TALE_MajSat_ strategy to specifically target DAXX, SETDB1 or SUV39H1 to chromocenters in Suv39dKO ESCs (Fig. 5E). Both histone methyltransferases were able to individually restore H3K9me3 at chromocenters, when targeted by TALE_MajSat_ (Fig. 5E; Fig S9C). The specific recruitment of DAXX to major satellite foci also promoted H3K9me3 deposition via SETDB1, at 70% of chromocenters. The impact of the different TALE_MajSat_ fusions on PCH clustering was further assessed by quantifying the number of TALE-bound foci (Fig. 5F). Consistent with our observations in *Daxx* KO and WT ESCs (Fig. 3D, S5A) targeting DAXX to major satellite also enhanced PCH clustering in Suv39dKO ESCs as evidenced by the reduced number of PCH foci (Fig. 5F). However, the recruitment of either SETDB1 or SUV39H1 did not affect the number of chromocenters, supporting the conclusion that H3K9me3 modification does not influence PCH clustering (Fig. 5F).

Overall, our results demonstrate that DAXX recruits a catalytically active SETDB1 to PCH upon 2iV conversion. However, SETDB1 and H3K9me3 modification alone are not sufficient to mediate PCH clustering, confirming that DAXX plays a central role in PCH organization in ground-state ESCs.

### H3.3K9 modification is crucial for PCH organization

Since DAXX, unlike SETDB1, had a direct impact on the spatial organization of chromocenters, we examined the role of H3.3 chaperone activity of DAXX by expressing a mutated form, DAXX^Y222A^, that lowers its interaction with H3.3 (Elsässer et al., 2012). We confirmed that co-immunoprecipitation of TALE_MajSat_ DAXX^Y222A^ fusion with H3.3 was drastically reduced compared to its wild-type counterpart (Fig. S9D). While targeting DAXX to chromocenters could recruit a tagged version of H3.3, neither the control TALE_MajSat_ nor TALE_MajSat_ DAXX^Y222A^ fusion were able to mobilize H3.3 to PCH (Fig. S9E; Fig. S9F). When expressed in Suv39dKO ESCs, the TALE_MajSat_ DAXX^Y222A^ fusion did not increase H3K9me3 levels at chromocenters, as it failed to recruit SETDB1 (Fig. 5E; Fig. S8C; Fig. S8D). Importantly, expressing the DAXX^Y222A^ mutant in *Daxx* KO ESCs did not change the number of chromocenters arguing that H3.3 is crucial for DAXX-mediated PCH clustering (Fig. 5F). To further confirm a function for H3.3 in chromocenter organization, we depleted endogenous H3.3 using a pool of siRNAs targeting the 3’UTR of H3f3a/b mRNAs and expressed TALE_MajSat_-DAXX in *Daxx* KO ESCs (Fig. 5G; Fig. S9E). The H3.3 knock-down was rescued by the expression of different HA-tagged versions of H3.3, which were assessed for their impact on PCH clustering by immuno-FISH (Fig. 5G, S9G). The binding of DAXX at major satellite did not enhance PCH clustering when H3.3 was knocked down. The expression of a wild-type H3.3-HA significantly reduced the number of major satellite foci and rescued the PCH hyperclustering phenotype mediated by TALE_MajSat_-DAXX. In contrast, expression of H3.3^G90M^, a mutant unable to bind DAXX (Elsässer et al., 2012; Hoelper et al., 2017), did not decrease the number of major satellite foci (Fig. 5H). This result is consistent with the absence of a chromocenter clustering phenotype in cells transfected by TALE_MajSat_-DAXX^Y222A^ (Fig. 5F) and confirms the important role of H3.3 in PCH clustering. Since DAXX can recruit SETDB1 to PCH, we asked whether a H3.3K9 mutant that cannot be methylated impacts PCH clustering and used H3.3^K9A^. Like H3.3^G90M^, H3.3^K9A^ failed to rescue the hyper-clustering phenotype in TALE_MajSat_-DAXX expressing cells, suggesting that the modification of this residue is essential for the spatial organization of chromocenters (Fig. 5H).

Altogether, we conclude that the role of DAXX in PCH organization is intrinsically linked to its chaperone activity. Interaction with H3.3 and H3.3K9 modifications are important in facilitating the spatial organization of pericentromeres in pluripotent ESCs.

## Discussion

During early embryogenesis, adapting heterochromatin to compensate for the wave of DNA demethylation is essential to maintain transcriptional repression of repetitive DNA and protect genome integrity. The mechanisms and molecular factors responsible for heterochromatin reorganization were, however, largely unknown. Here, we describe a novel and essential role for the H3.3-chaperone DAXX and its partner PML in the survival of ground-state pluripotent stem cells. Taken together, our results support a model in which DAXX and PML facilitate both pericentric and peripheral heterochromatin formation in ground-state ESCs (Fig. 6). At PCH, DAXX deposits H3.3 and recruits PML and SETDB1, which can methylate H3.3K9. In the absence of either DAXX or PML, PCH is compromised (Fig. 6). In *Daxx* KO ESCs, heterochromatin at major satellite is less compact and partially loses its boundary properties leading to defective chromocenter clustering. At the nuclear periphery, PML and DAXX are necessary for the maintenance of the heterochromatin positioned beneath the nuclear envelope, contributing to the transcriptional silencing of genes tethered to the nuclear lamina. The global failure in heterochromatin compartment formation could directly contribute to the impossibility of *Daxx* KO and *Pml* KO ESCs to maintain the ground-state of pluripotency.

**Fig. 6.**
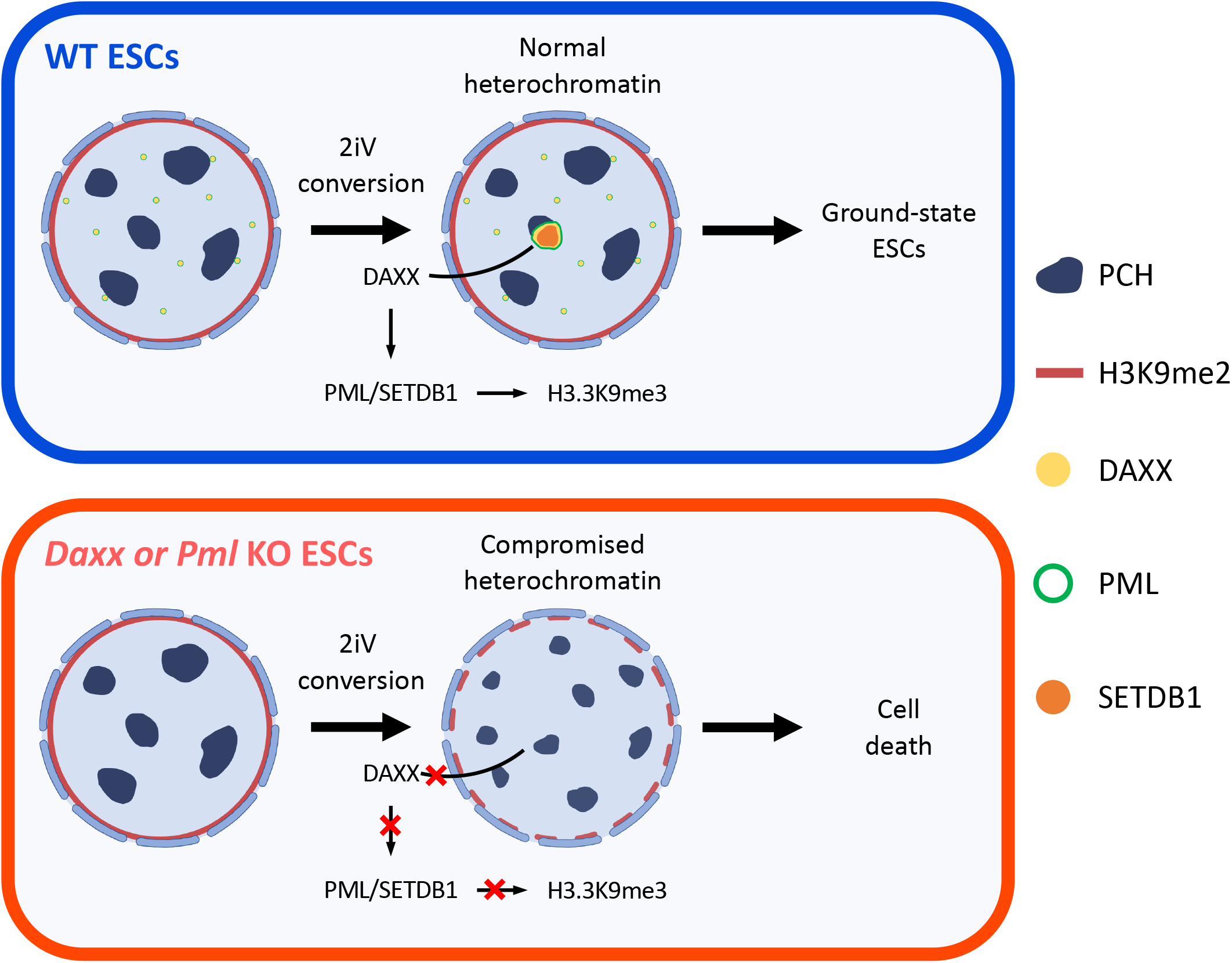
Model for heterochromatin maintenance by DAXX in ESCs. During ground-state conversion of ESCs, DAXX accumulates at PCH and is surrounded by a PML shell. DAXX recruits the histone methyltransferase SETDB1 to promote H3.3K9me3 to maintain the pericentromeric heterochromatin. In *Daxx* and *Pml* KO cells, the 3D-organization of heterochromatin is impaired. The peripheral heterochromatin H3K9me2 fails to accumulate properly at the nuclear edge leading to promiscuous expression of genes located in LADs. The clustering and the physical properties of pericentromeric heterochromatin is impacted, impairing transcriptional repression of major satellites. The global defect of heterochromatin observed in these cells prevents prolonged culture of these cells in ground-state condition.

### DAXX and H3.3 impact chromocenter organization

Both DAXX and H3.3 are essential for early embryogenesis, but their specific functions have remained elusive (Jang et al., 2015; Lin et al., 2013; Liu et al., 2020; Michaelson et al., 1999). we found that DAXX is necessary for maintaining the spatial organization and chromatin state of pericentric heterochromatin (PCH), thereby protecting the viability of ground-state ESCs (Fig. 3B; Fig. 3F; Fig. 3H). This finding is consistent with *in vivo* observations, which show that DAXX and SETDB1 are recruited to PCH just before chromocenter formation, and the viability of *Daxx* KO embryos drops quickly after the transition to the pluripotent stage (Arakawa et al., 2015; Cho et al., 2012; Liu et al., 2020). These observations support our conclusion that the role of DAXX and SETDB1 at PCH might be crucial for the ground-state of pluripotency. Since defective PCH assembly is often associated with genomic instability, our model may also explain why *H3f3a/b* double knockout ESCs exhibit severe chromosome segregation defects (Jang et al., 2015).

While *Daxx* deletion had no impact on major satellite transcription and chromocenter formation upon neuronal differentiation (Fig. 2A; Fig. 3B), DAXX and H3.3 are important for chromocenter clustering during myoblast differentiation (Park et al., 2018; Salsman et al., 2017). In myoblasts, DAXX is recruited to chromocenters, where H3.3 deposition stimulates transcription of major satellites, suggesting a mechanism different to the one we observed in ESCs. Different DAXX and partner recruitment pathways could explain the opposite impact of DAXX and H3.3 on major satellite transcription. In ESCs, DAXX recruits SETDB1, thus favoring a repressive chromatin environment. Upon myoblast differentiation, *Setdb1* is downregulated and might not be available for interaction with DAXX, which is recruited to major satellite repeats by the muscle-specific lncRNA ChRO1 (Park et al., 2018; Song et al., 2015).

Recent insights into chromocenter organization have come from the observation that the intrinsically disordered regions of HP1α facilitate the compartmentalization of chromocenters through liquid-liquid phase separation (Larson et al., 2017; Strom et al., 2017). Since the first fifty amino-acids and the C-terminal half of DAXX are intrinsically disordered (Escobar-Cabrera et al., 2010), it is possible that DAXX may also undergo liquid-liquid phase separation. Multivalent interactions between chromodomains recognizing H3K9me3 have also been proposed to contribute to phase separation of heterochromatin (Wang et al., 2019). Yet, *Daxx* deletion impacted the boundary properties of pericentromeric heterochromatin without affecting H3K9me3 (Fig. 3H; Fig. S9A). Since DAXX-mediated chromocenter clustering relies on H3.3K9 modification (Fig. 5H), future work will compare the interaction of HP1α and additional heterochromatin-related proteins to H3.3K9me3 vs. H3K9me3.

### Mechanism of DAXX targeting to pericentric heterochromatin

In ground-state ESCs, DAXX binds only a subset of chromocenters in approximatively 80% of the population (Fig. 2E). Given that chromocenters replicate synchronously during mid-S phase (Guenatri et al., 2004), the recruitment of DAXX is unlikely to rely on replicative chromatin remodeling. DNA hypomethylation could be one of the factors responsible for DAXX mobilization to PCH. Polycomb group proteins have been shown to be recruited to PCH specifically under hypomethylated conditions (Pailles et al., 2022; Saksouk et al., 2014; Tosolini et al., 2018). The fact that DAXX was shown to rely on PRC1 to bind major satellites in zygotes would support a polycomb-mediated recruitment of DAXX in ground-state ESCs (Liu et al., 2020). However, the presence of H3K27me3, deposited by PRC2, is relatively homogenous amongst the PCH foci within a nucleus, suggesting a different mechanism (Tosolini et al., 2018).

DNA damage may also be a factor in the recruitment of DAXX to PCH. Several studies have described the incorporation of H3.3 after DNA damage (Adam et al., 2013; Fortuny et al., 2021; Juhász et al., 2018; Li and Tyler, 2016; Luijsterburg et al., 2016). Both HIRA and DAXX-ATRX complexes have been proposed to deposit H3.3 after DNA damage, but DAXX has been more specifically observed at chromocenters damaged by UVC (Fortuny et al., 2021). Active DNA demethylation may be a source of DNA damage responsible for DAXX recruitment. DAXX relocation to PCH was specific to the 2iV culture condition (Fig. 2B), a medium that was shown to stimulate TET enzymes activity (Blaschke et al., 2013). *In vivo,* DAXX binds preferentially paternal chromosomes which are actively demethylated by the TET enzymes, a pathway that can result ultimately in a DSB (Nakatani et al., 2015; Wossidlo et al., 2011).

### Role of SETDB1 at chromocenters in ground-state ESCs

SETDB1 has been shown to be responsible for H3K9me3 deposition at transposable elements and telomeres (Elsässer et al., 2015; Gauchier et al., 2019; Karimi et al., 2011). Yet, knocking down SETB1 in Suv39h1/h2 double knock-out cells destabilizes chromocenters suggesting that SETDB1 could be involved in PCH formation (Pinheiro et al., 2012). Our data show that the chaperone activity of DAXX is necessary to recruit SETDB1 to pericentromeric heterochromatin (Fig. S8C). In contrast to SUV39H1/H2, SETDB1 contains a triple Tudor domain recognizing the double modification K14 acetylation and K9 methylation that may facilitate its binding to hyperacetylated, newly incorporated histone H3.3 (Jurkowska et al., 2017). Using ESCs lacking Suv39h1/h2, we found that SETDB1 can deposit H3K9me3 at chromocenters (Fig. 5C; Fig. 5D). This function of SETDB1 is consistent with its role at transposable element in ESCs (Elsässer et al., 2015; Hoelper et al., 2017). Yet, it contrasts with the observation that SETDB1 promotes H3K9me1 at major satellites in mouse embryonic fibroblasts (Loyola et al., 2009). However, SETDB1 was specifically recruited during S-phase by the H3.1/H3.2 chaperone, CAF1, suggesting that SETDB1 substrate specificity might change depending on its recruitment pathway. Similar to our data in ESCs, CAF1 recruited SETDB1 to chromocenters only in a subset of S-phase cells. Since pericentromeric satellites were shown to be particularly sensitive to replicative stress, SETDB1 recruitment could also result from DNA damage (Crosetto et al., 2013).

### The role of DAXX and H3.3 upon DNA hypomethylation

This study reveals that DAXX is recruited to pericentromeres in the context of DNA hypomethylation. While PML nuclear bodies are generally devoid of chromatin, PML formed a shell around DAXX-positive chromocenters in hypomethylated ground-state ESCs (Fig. 4A). Similar structures have been observed in patients with immunodeficiency, centromeric instability and facial dysmorphia (ICF) syndrome associated with mutations in DNA methyltransferases (Luciani et al., 2006). In patient lymphocytes, DAXX and DNA repair proteins accumulate at hypomethylated pericentromeric satellites, suggesting that our proposed model could apply to other pathologies (Fig. 6).

In conclusion, our study reveals DAXX as an important factor for different heterochromatic compartments in ground-state ESCs. At the nuclear periphery, DAXX and its partner PML are necessary for the clustering of peripheral heterochromatin at the nuclear edge. At PCH, DAXX deposits H3.3 and recruits PML and SETDB1 to facilitate chromocenter formation by enhancing their clustering properties. It would be interesting to characterize whether DAXX also contributes to the 3D-organization of other heterochromatin domains, such as the clustering of LINE1 elements (Lu et al., 2021). Beyond early development, our work could also provide critical insights into the molecular pathways that overcome DNA hypomethylation transitions in pathological contexts, such as cancers.

## Material and Methods

### Cell lines

Feeder-free E14 mES cells were used for most experiment excepts otherwise notified. E14 mES cells were kindly provided by Pablo Navaro (Pasteur Institute). *Daxx* KO cell line was constructed using CRISPR/cas9 editing in E14 mES cells. Guide-RNA was designed using the online CRISPOR tool. Oligos were designed with a BbsI site on 5’ to clone them into the pSpCas9(BB)-2A-Puro(pX459) v2.0 vector (Ran et al., 2013). Guide was designed carrying two guanines on 5’ of the sequence to avoid off-target effects, as previously described (S. W. Cho et al., 2014). Sg-mDaxx: gGACCTCATCCAGCCGGTTCA. *Pml* KO ESCs were generated by CRISPR from E14 cells and described in a previous study (Tessier et al., 2022). Feeder-free Suv39HdKO and corresponding WT R1 ES cells were provided by Alice Jouneau and generated in Antoine Peters’s laboratory (Lehnertz et al., 2003).

### Culture conditions

Pluripotent cells were cultured either in a serum condition, defined as follow: DMEM (Gibco), supplemented with 10% of ESC-certified FBS (Gibco), 2-mercaptoethanol (0.05mM, Gibco), Glutamax (Gibco), MEM non-essential amino acids (0.1mM, Gibco), Penn-Strep (100units/mL, Gibco) and LIF (1000units/mL,) for serum condition. Serum-cultivated cells were grown on 0.1% gelatin-coated plates or stem cell plates (Stem Cell technology) at 37°C with 5% CO2. Medium was changed every day and cells were passaged every 2 to 3 days. The other culture condition is the chemically defined serum-free 2i condition defined as follow: Neurobasal:DMEM/F-12 (50:50, Gibco) medium, supplemented with N2 and B-27 supplements (Gibco), BSA fraction V (0.05%Gibco), 1-thioglycerol (1.5×10^-4^M, Sigma) and ESGRO 2 inhibitors (GSK3i and MEKi) and LIF (Merck). Vitamin C (L-ascorbic acid 2-phosphate, Sigma) was added at a concentration of 100μg/mL (Blaschke et al., 2013). Cells were grown on 0.1% gelatin-coated plates. Medium was changed daily. Cells were passed every 2 days at 1:4 ratio for the first passage then at a 1:6 ratio. Differentiation of mES cells was done by LIF removal for the first 24h. Then, non-LIF medium was supplemented with retinoic acid (10^-6^M,) for 4 days. For cell growth quantification, cells were counted at each passage for at least 2 different experiments.

### Vectors and transfections

Cells were harvested using trypsin and one million was plated in a 0.1% gelatin-coated plate and transected with 0.2 to 2.5μg of DNA using the Lipofectamine 2000 reagent (Thermo), following manufacturer’s protocol. TALE vectors were constructed using previously described methods (Ding et al., 2013). A TALE specific DNA binding domain targeting *Major Satellite* repeats was created by the modular assembly of individual TALE repeats inserted into a backbone vector containing TALE-Nrp1-VP64 previously described (Therizols et al., 2014). The BamHI-NheI fragment containing VP64 was replaced by PCR products encoding the DAXX protein, corresponding DAXX mutants or additional proteins such as SUV39H1 and TET1CD. CDS of the different proteins were amplified by PCR from cDNAs obtained after RNA extraction of serum E14 mES cells. DAXX mutants were generated by PCR from WT DAXX.

Oligos for major satellites targeting were designed with a BbsI site on 5’ to clone them into the pSpCas9(BB)-2A-Puro(pX459) v2.0 vector (Ran et al., 2013). Catalytically dead Cas9 (dCas9) was inserted into the pSpCas9-2A-Puro(pX459) v2.0 to generate a SpdCas9-2A-Puro.

Target sequences for the TALE repeat domains and dCas9 associated guides RNA are listed in supplementary table.

### RNA extraction for RNAseq or RT-qPCR

Total RNA was extracted using RNeasy extraction kit (Qiagen) according with manufacturer’s protocol including DNAseI treatment for 15min at room temperature (Qiagen). Complementary DNA were generated from 1μg of RNA using the Maxima first strand cDNA synthesis kit (Thermo Fisher), with a second round of DNAseI from the Maxima kit for 15min. Real-time qPCR was carried out using a LightCycler 480 instrument (Roche) and the LightCycler 480 SYBR green master mix (Roche). The qRT-PCR primers used in this study are listed in supplementary table. Three independent biological repeats were obtained for each sample. For RNAseq experiment, RNA quality was assessed using the Agilent 2100 bioanalyzer. Libraries were prepared using oligo(dT) beads for mRNA enrichment, then fragmented and reverse transcribed using random hexamers primer. After adaptor ligation, the double-stranded cDNA is completed through size selection of 250-300bp and PCR amplification, then quality of the library is assessed by the Agilent 2100 bioanalyzer. Sequencing was performed in 150bp paired-end reads using an Illumina sequencer platform.

### RNA-seq Mapping and Processing

FASTQ files generated by paired end sequencing were aligned to the mouse genome using bowtie2 v2.2.6 (parameters: --local --threads 3; mm9 genome build). Mapped RNA-seq data was processed using tools from the HOMER suite (v4.8). SAM files were converted into tag directories using ‘makeTagDirectory’ (parameters: -format sam -sspe). Genomic intervals which extended beyond the end of the chromosomes was removed using ‘removeOutOfBoundsReads.pl’. bigWig browser track files were generated using ‘makeUCSCfile’ (parameters: -fsize 1e20 -strand + -norm 1e8). For gene expression analysis, read depths were quantified for all annotated refseq genes using analyzeRepeats.pl (parameters: rna mm9 -strand both -count exons -rpkm -normMatrix 1e7 -condenseGenes).

For repeat analysis, read coverage was quantified for each repeat and then condensed to a single value for each named entry (parameters: repeats mm9 -strand both -rpkm -normMatrix 1e7 -condenseL1). Read depths were then corrected for the number of instances of each repeat prior to expression analysis.

### Expression Analysis

Quantified RNA-seq data was processed using the limma package (R/Bioconductor)(Team, 2017). Following the addition of an offset value (1 RPKM) to each gene or repeat, data was normalised across all samples using ‘normalizeBetweenArrays’ with method=‘quantile’. Foldchanges and p-values for differential expression of genes and repeats were determined using empirical Bayes statistics. Briefly, data was fit to a linear model using ‘lmFit’ and specified contrasts were applied using ‘makeContrasts’ and ‘contrasts.fit’. Data was processed using the ‘topTable’ function with adjust.method=“BH” (Benjamini-Hochberg multiple-testing correction). Differential expression was defined as log2 fold change ≤-1 or ≥ 1 and an adjusted p-value of ≤ 0.01. Three biological replicates for each condition represent independently cultured pools of cells.

### Data visualization

Heatmaps and boxplots were generated using Prism GraphPad (v9). Histograms were drawn using either Prism GraphPad or Excel. Volcano plots were generated using the plot function in R.

### Immunofluorescence

Murine ES cells were harvested with trypsin (Gibco) and plated for 4-12h onto either 0.1% gelatin-coated or ECMatrix-coated (Sigma) glass cover slips. Cells were fixed with 4% paraformaldehyde for 10min at room temperature, then rinsed three times with PBS. Cells were permeabilized with 0.1X triton for 12min at room temperature, then rinsed three times with PBS. Blocking was done in 3% BSA solution for 30min at room temperature. All incubations with primary antibodies were performed for either 1h at room temperature or overnight at 4°C with the following antibodies for H3K9me3 (Active Motif, 1:1000), DAXX (Santa Cruz, 1:500), H3K9me2 (Active Motif, 1:1000), LaminB1 (Abcam, 1:1000), PML () and SETDB1 (Proteintech, 1:100). Incubation with secondary antibodies (fluorescently labeled anti-mouse or anti-rabbit, 1:1000) were performed for 1h at room temperature. Mounting was performed using ProLong Diamond with DAPI mounting media (Thermo). Antibodies are listed in supplementary table.

### Fluorescent in situ Hybridization

Murine ES cells were harvested with trypsin (Gibco) and plated for 4-6h onto 0.1% gelatin-coated glass cover slips. Cells were fixed with 4% paraformaldehyde (PFA) for 10min at room temperature, then rinsed three times with PBS. Cells were permeabilized with 0.5X triton for 12min at room temperature, then rinsed three times with PBS. Cells were briefly washed in 2X SSC, then treated with RNAseA (100μg/mL, Sigma) for 1h at 37°C. Cells were briefly washed in 2X SSC, then denatured by serial 2min incubation into 70,90 and 100% ethanol. Cover slips were air dried for 15min. Cover slips are incubated with 200nM of PNA probe, placed for 10min at 95°C for denaturation, then placed for 1h at room temperature in the dark for hybridization. Cover slips were washed twice in 2X SSC 0.1%Tween-20 for 10min at 60°C. Cover slips were immerged at room temperature in 2X SSC 0.1%Tween-20 for 2 min, then in 2X SSC for 2min and 1X SSC for 2min. Mounting is performed using ProLong Diamond with DAPI mounting media (Thermo).

### Dot blot experiments

DNA extraction was performed using the Wizard genomic DNA extraction kit (Promega). Genomic DNA (1μg) was then denatured in 0.1M NaOH for 10min at 95°C before neutralization with 1M NH4OAc on ice for 5min. DNA samples were spotted on a nitrocellulose membrane. Blotted membrane was washed in 2X SSC and dried at 80°C for 5min before UV cross-linking at 120,000μJ/cm2. Membrane was then blocked using PBS, 5% milk 0.1% tween for 30min at room temperature. Membrane was incubated with 5mC antibody (Diagenode, 1:1000) for 3h at room temperature. After 3 washes of 10min each in PBS, membrane was blocked again for 30min and then incubated with secondary anti-HRP antibody for 1h at room temperature. Membrane was washed 3 times for 10min in PBS and visualized by chemiluminescence with ECL Plus.

### Fluorescent Recovery After Photobleaching and variance analysis

Murine ES cells transfected with HP1α-GFP were harvested with trypsin (Gibco) and plated for 4-6h onto 0.1% gelatin-coated glass live-cell Nunc slides (Thermo). FRAP experiment was carried with an LSM 800 confocal microscope (Zeiss). The 488nm laser was used to bleach and acquire GFP signal. 1 image was taken before a bleach pulse of 5ms. Bleaching area was set to target a single pericentric domain. Images were acquired every second during 35s post-bleach. FRAP analysis was performed using a FRAP analysis ImageJ Jython script, that generated FRAP curves and the associated half-recovery time and mobile fraction parameters. GFP-HP1α variance along time was obtained from ImageJ analysis using standard deviation z-projection along time for the whole duration of the movie. Quantification of heterochromatin barriers were performed using a 1μm line across individual non-bleached chromocenter borders, for which the variance intensity along time was measured with the ImageJ software.

### Image acquisition and analysis

Images for immunofluorescence and FISH experiments were obtained with an inverted Nikon Ti Eclipse widefield microscope using a 60X immersion objective and LED sources. Z-stacks images were taken and then deconvoluted using a custom ImageJ deconvolution script. Quantifications of images were performed using custom Icy scripts in Icy and ImageJ. To quantify DAXX enrichment at PCH and the number of PCH foci, chromocenters were segmented using ICY the spot detector function on either H3K9me3 or major satellite FISH signal. Nuclei were isolated using the ICY k-means segmentation script. The nuclear periphery enrichment was measured in ImageJ, briefly linescans of LaminB1 and H3K9me2 at the periphery were isolated using the straighten command and subsequently used to generated intensity profiles.

H3K9me3 intensities at chromocenters were measured using à 2μm line across individual DAPI-dense chromocenters. Two to three chromocenters per nucleus were analyzed.

### Statistical analysis

Number of objects counted, and statistical tests performed are indicated in the text, figure or figure legends. All statistical analysis results are listed in supplementary file. Pvalues are represented as follow: * <0.05; **<0.01; ***<0.001; ****<0.0001.

### Western Blotting

Total protein extracts were prepared in RIPA buffer with protease inhibitor cocktail (Roche). Samples were sonicated for 3 minutes alterning 30seconds ON/ 30seconds OFF. Proteins were separated by electrophoresis in 8-15% poly-acrylamide gels then transferred onto nitrocellulose membranes. Membranes were incubated in ponceau then washed in PBS, 0.1% tween. Membranes were blocked in PBS, 0.1% tween, 5% milk for 30min at room temperature before incubation with primary antibody overnight at 4°C. After 3 washed in PBS, 0.1% tween, membranes were incubated with secondary HRP-conjugated antibody for 1h at room temperature. Membranes were washed 3 times in PBS, 0.1% tween and visualized by chemoluminescence using ECL Plus.

## Acknowledgements

We would like to thank A. Bonnet-Garnier and A. Jouneau for the gift of Suv39dKO KO ESCs and advice. We are also grateful to image core microscopy facilities of IRSL, Paris, in particular N. Setterblad for his assistance with live imaging. We warmly thank V. Lallemand-Breitenbach and H. de Thé (Collège de France, Paris, France) for the sharing of PML antibody and helpful advice. We are thankful to Y. Khalil for her help in microscopy analysis. We kindly thank P. Lesage and Annia Carré Simon for their critical readings of the manuscript, and other members of the team for helpful advice. We finally thank support services of IRSL. This work was funded by Agence Nationale pour la Recherche (ANR-16-CE12-0003), IDEX SLI, La Ligue Nationale Contre le Cancer (LNCC)-Comité Ile de France and the labex Who am I. AC was supported by MESR and LNCC. The authors declare no conflict of interest.

## Author Contributions

AC, AV, CD, PT, performed all experiments, RI analyzed RNAseq data. AC, AV, PT designed experiments, interpreted data. RB wrote the scripts for image analysis. AC, EF, PT contributed to the writing of the manuscript. All authors reviewed the manuscript.

## Declaration of interest

The authors declare no competing interests.

## Supplemental material legends

**Fig. S1 | Impact of *Daxx* deletion on ground-state conversion. (A)** Representative field of WT and *Daxx* KO ESCs after IF against DAXX (green) and counterstaining with DAPI (cyan). Scale bar = 5μm. **(B)** Dot blot using 5mC antibody to assess global DNA methylation levels upon 2iV conversion. One experiment is shown. Histogram displays mean with SEM from 3 independent replicates. **(C)** Volcano plot representing the number of differentially expressed genes in ground-state, serum or differentiated cells from RNA-seq experiments as in C. Differentially expressed genes in red for down-regulated or blue for up-regulated correspond to Log2(FoldChange(KO/WT)) <-1 or >1 with p-value<0.05. **(D)** Heatmaps representing genes displaying strongest transcriptional changes upon 2iV conversion (left) or upon differentiation (right) normalized by transcription in serum conditions. Same set of genes for the *Daxx* KO ESCs. Data are from 3 biological replicates of RNA-seq experiments. **(E)** Volcano plots representing the number of differentially expressed genes upon 2iV conversion of WT and *Daxx* KO ESCs. Previously identified 2iV conversion markers are in red for up-regulated or blue for down-regulated (Blaschke et al., 2013).

**Fig. S2 | Impact of *Daxx* deletion on nuclear periphery. (A)** RT-qPCR RNA quantification LAD genes in WT and *Daxx* KO ESCs in serum and upon 2iV conversion. Histogram represents mean normalized to WT-serum with SEM of 3 biological replicates. **(B)** Left, representative image of ESCs after IF against H3K9me2. Top right, linescan represents straighten H3K9me2 signal at the nuclear periphery (black). Bottom right, Intensity profiles of IF signals measured on linescans at the periphery (blue) or in the center (orange) of the nucleus. Ratio between mean intensity at the periphery and in the center was used to determined peripheral enrichment. **(C)** Representative nuclei after immuno-detection of LaminB1 (red) and counterstaining with DAPI (blue) in WT and *Daxx* KO ESCs in serum and 2iV (scale bar = 2μm). Linescan represent straighten LaminB1 signal at the nuclear periphery (black). **(D)** Boxplots of LaminB1 signal quantification enrichment at the nuclear periphery from 2 independent replicates. Mann-Whitney-tests were used for statistical analysis.

**Fig. S3 | Impact of *Daxx* deletion on DNA repeats silencing.** RT-qPCR RNA quantification of DNA repeats. L1Tf, IAP elements and major satellites in WT and *Daxx* KO ESCs in serum, and upon differentiation or 2i+VitC (2iV) conversion. Histogram represents mean with SEM of 3 biological replicates.

**Fig. S4 | Quantification of DAXX at chromocenters. (A)** Representative immunoFISH picture with H3K9me3 and major satellites DNA. **(B)** Dot plots of Pearson correlation between H3K9me3 and major satellite signals. **(C)** Representative field of ESCs grown in serum or 2iV immuno-detection of H3K9me3 (red), DAXX (green) and counterstaining with DAPI (cyan). White squares and arrows highlight the presence (2iV) of larger DAXX foci with higher levels of H3K9me3 and DAPI. Scale bar = 10μm. **(D)** Distribution of DAXX foci volume observed in WT ESCs grown in serum (black) or in 2iV (red) from 3 independent replicates. Dashed line indicates the threshold used to defined large or small DAXX foci. **(E)** Boxplots of DAPI signal intensity observed under small and large DAXX foci in serum (white) and 2iV (red) conditions. **(F)** Boxplots of H3K9me3 signal intensity observed under small and large DAXX foci in serum (white) and 2iV (red) conditions. **(G)** Top, ESCs nucleus after immuno-detection of H3K9me3, DAXX and counterstaining with DAPI. Bottom, for each nucleus, H3K9me3 and DAPI signals were segmented to PCH and nuclear ROIs. DAXX enrichment at PCH was defined by yhe ratio of DAXX intensity under PCH ROIs. Mann-Whitney-tests were used for statistical analysis in B, E and F.

**Fig. S5 | Role of DAXX on pericentromeric heterochromatin formation. (A)** Left, representative immunoFISH pictures for major satellites and Flag for WT ESCs, grown in serum, transfected with TALE_MajSat_-Δ or TALE_MajSat_-DAXX. Right, quantification of the number of major satellite foci detected in the focal plane and corresponding surface area. n=total number of nuclei for number and foci for sizes analyzed. Two-sided Mann-Whitney tests were used for statistical analysis. **(B)** Left, mobile fractions of GFP-HP1α calculated from data presented in Fig. 4D. n=total number of cells analyzed from 2 biological replicates. Right, fluorescence recovery curves generated for FRAP experiments in WT or *Daxx* KO cells in serum or ground-state condition. **(C)** MNAse digestion for indicated times in minutes (min.) in WT or *Daxx* KO cell lines. Total DNA is revealed on a BET agarose gel and Major satellites via a Southern blot experiment. Right, intensity values of total DNA or major satellite signals, according to fragment size, from 4min of MNase digestion. One experiment is shown from 2 biological replicates.

**Fig. S6 | Quantification of PML at chromocenters. (A)** Representative field of ESCs grown in serum or 2iV after immuno-detection of H3K9me3 (red), PML (green) and counterstaining with DAPI (cyan). Scale bar = 10μm. **(B)** Distribution of PML foci volume observed in WT ESCs grown in serum (black) or in 2iV (red) from 3 independent replicates. Dashed line indicates the threshold used to defined large PML foci. **(C)** Boxplots of DAPI signals intensities observed under small and large PML foci in serum (white) and 2iV (red) conditions. **(D)** Boxplot of H3K9me3 signals intensities observed under small and large PML foci in serum (white) and 2iV (red) conditions. **(E)** Representative nucleus of ESCs grown in 2iV after immuno-detection of H3K9me3 (red), PML (green) and counterstaining with DAPI (cyan) presenting a PML-arc around a PCH cluster (white arrow). Scale bar = 2μm. Two-sided Mann-Whitney tests were used for statistical analysis in B and D.

**Fig. S7 | Impact of *Pml* deletion on nuclear periphery. (A)** Representative field of WT, *Daxx* KO and *Pml* KO ESCs, in serum or 2iV after IF against PML (green) and counterstaining with DAPI (cyan). **(B)** Representative nuclei after immuno-detection of LaminB1 (red) and counterstaining with DAPI (blue) in WT and *Pml* KO ESCs in serum and 2iV (scale bar = 2μm). Linescans represent straighten LaminB1 signal at the nuclear periphery (black). **(C)** Boxplots of LaminB1 signal quantification enrichment at the nuclear periphery from 2 independent replicates. WT values are taken from Fig. S2D. Two-sided Mann-Whitney tests were used for statistical analysis.

**Fig. S8 | DAXX recruits SETDB1 to pericentromeres. (A)** Representative immunofluorescence pictures for SETDB1, DAXX and PML in WT or *Daxx* KO ESCs in serum condition. **(B)** Top, representative immunofluorescence pictures for SETDB1, DAXX and PML in WT ESCs in serum. (**C)** Representative immunofluorescence pictures for SETDB1 and Flag-TALE in *Daxx* KO serum ESCs transfected with TALE_MajSat_ (tMs-Δ), TALE_MajSat_-DAXX (tMs-DAXX) or TALE_MajSat_-DAXX^Y222A^ (tMs-DAXXY222A). White dashed squares highlight DAPI-dense Flag-positive chromocenters. **(D)** Quantification of SETDB1 recruitment to Flag-positive chromocenters. Histogram represents mean with SEM from at least 2 biological replicates. n=total number of nuclei analyzed. Chi-square tests were used for statistical analysis.

**Fig. S9 | Role of H3.3 and H3K9me3 modifications at PCH. (A)** Representative immunoFISH pictures for H3K9me3 and DNA FISH of major satellites (MajSat) in both WT and *Daxx* KO ESCs in serum or ground-state (2iV) conditions. **(B)** Mean and SEM of fluorescence intensity of H3K9me3 measured over major satellite signal. n=total number of nuclei analyzed from two independent replicates. **(C)** Quantification of the percentage of cells displaying H3K9me3 recruitment at Flag-DAPI-dense PCH in Suv39dKO serum ESCs transfected with TALE_MajSat_-Δ, TALE_MajSat_-SUV39H1, TALE_MajSat_-SETDB1, TALE_MajSat_-DAXX or TALE_MajSat_-DAXX^Y222A^ grown in serum transfected with TALE_MajSat_-Δ, TALE_MajSat_-DAXX or TALE_MajSat_-DAXXY222A. T-tests were used for statistical analysis. **(D)** Represents signal after Flag-TALE immunoprecipitation. Bottom line represents input line from total extract. **(E)** Representative immunofluorescence pictures of HA-H3.3 in WT cells grown in serum transfected with TALE_MajSat_-Δ, TALE_MajSat_-DAXX or TALE_MajSat_-DAXX^Y222A^. **(F)** quantification of H3.3 recruitment at DAPI-dense chromocenters in the indicated Tale-transfected cells. Histogram represent mean with SEM from at least 2 biological replicates. n=total number of nuclei analyzed. T-tests were used for statistical analysis. **(G)** Western blot showing the amount of TALE and H3.3 variants in *Daxx* KO ESCs grown in serum transfected with the indicated constructions.

## Notes

### Competing Interest Statement

The authors have declared no competing interest.

### Summary of Updates

Section on image analysis has been redone with automated script ; Added a section on nuclear periphery ; Added datas on PmlKO ESCs ; In consequence all figure have been revised; author affiliations updated; Supplemental files updated.

