## Supplemental Figure for "DAXX safeguards heterochromatin formation in embryonic stem cells"

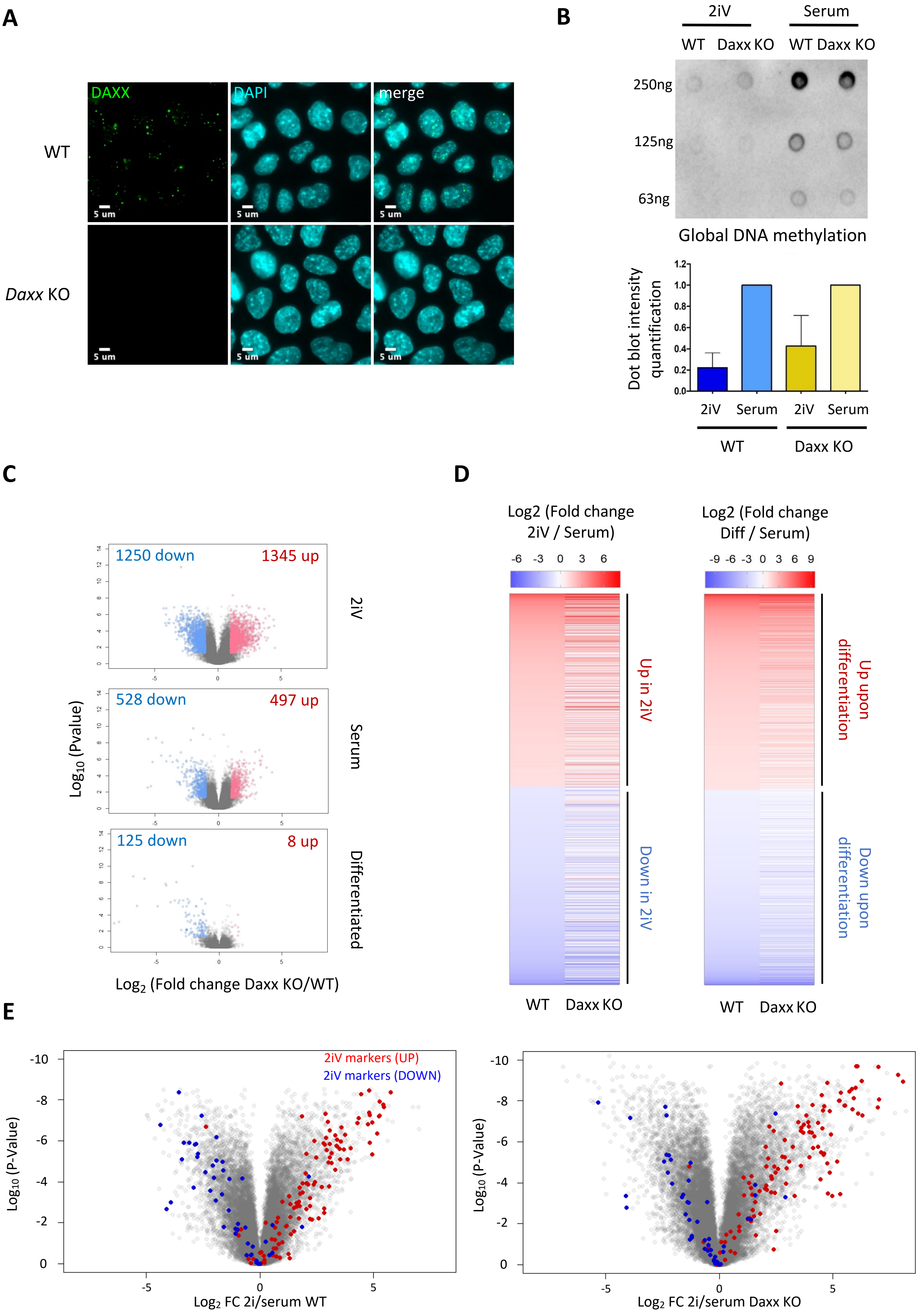


Figure S1. Canat et al. 2023

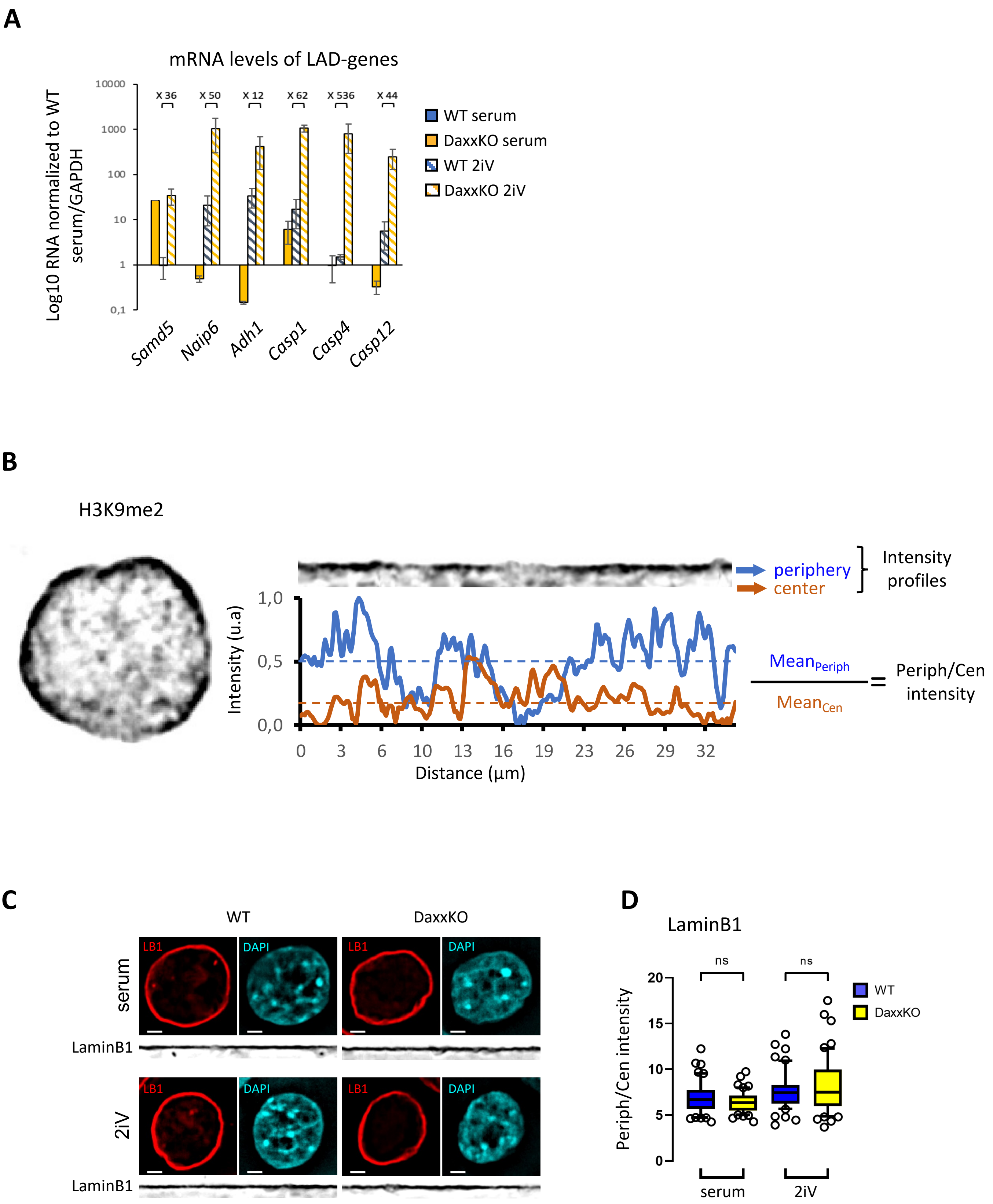


Supplemental Figure S2. Canat et al. 2023

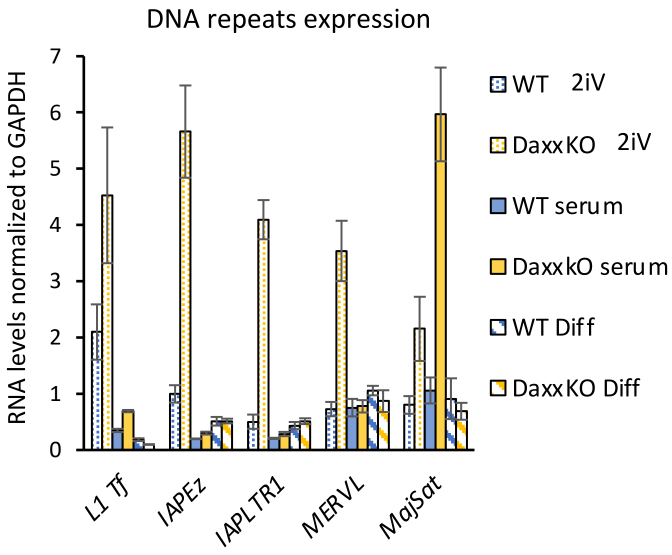


Supplemental Figure S3. Canat et al. 2023

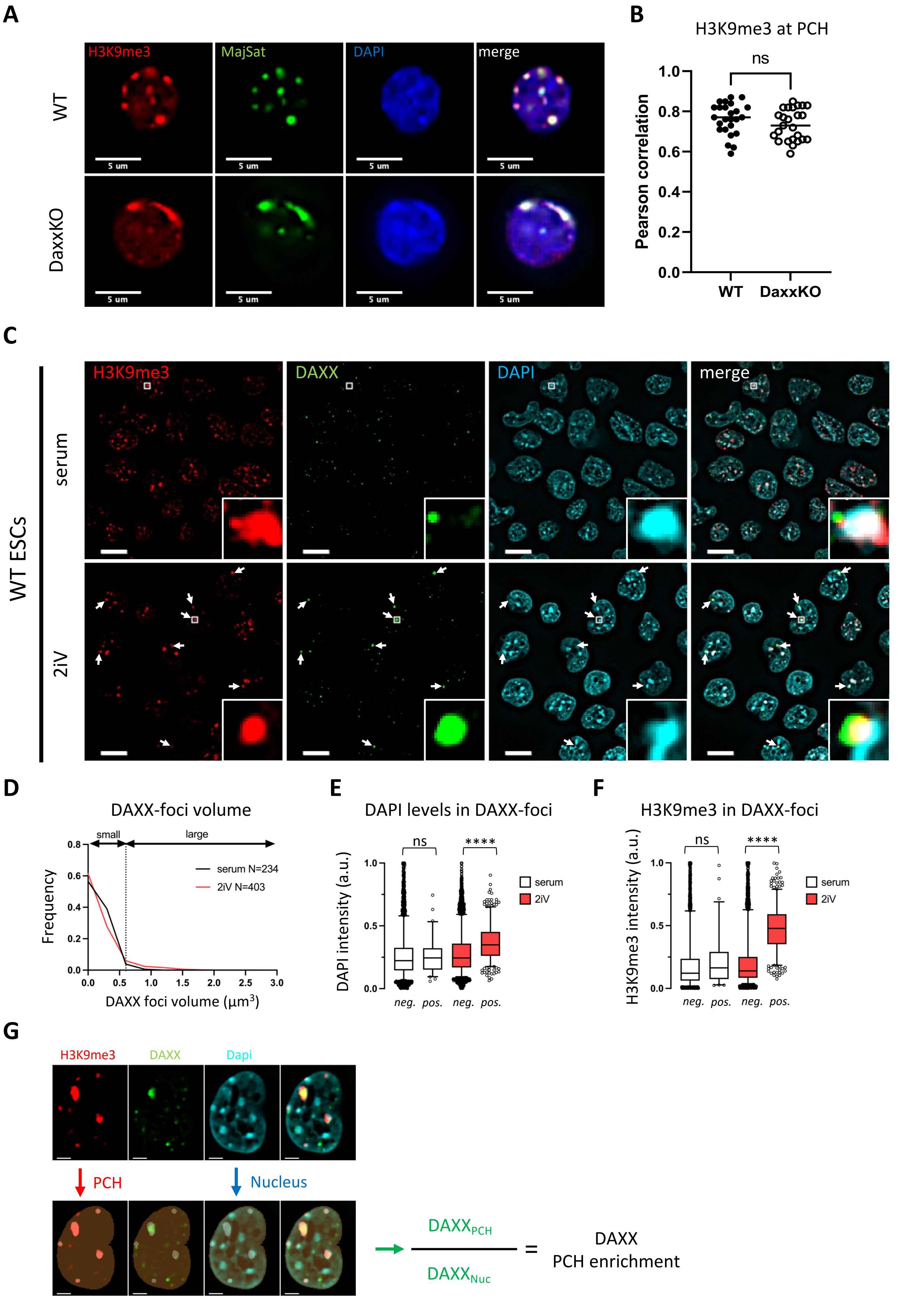


Supplemental Figure S4. Canat et al. 2023

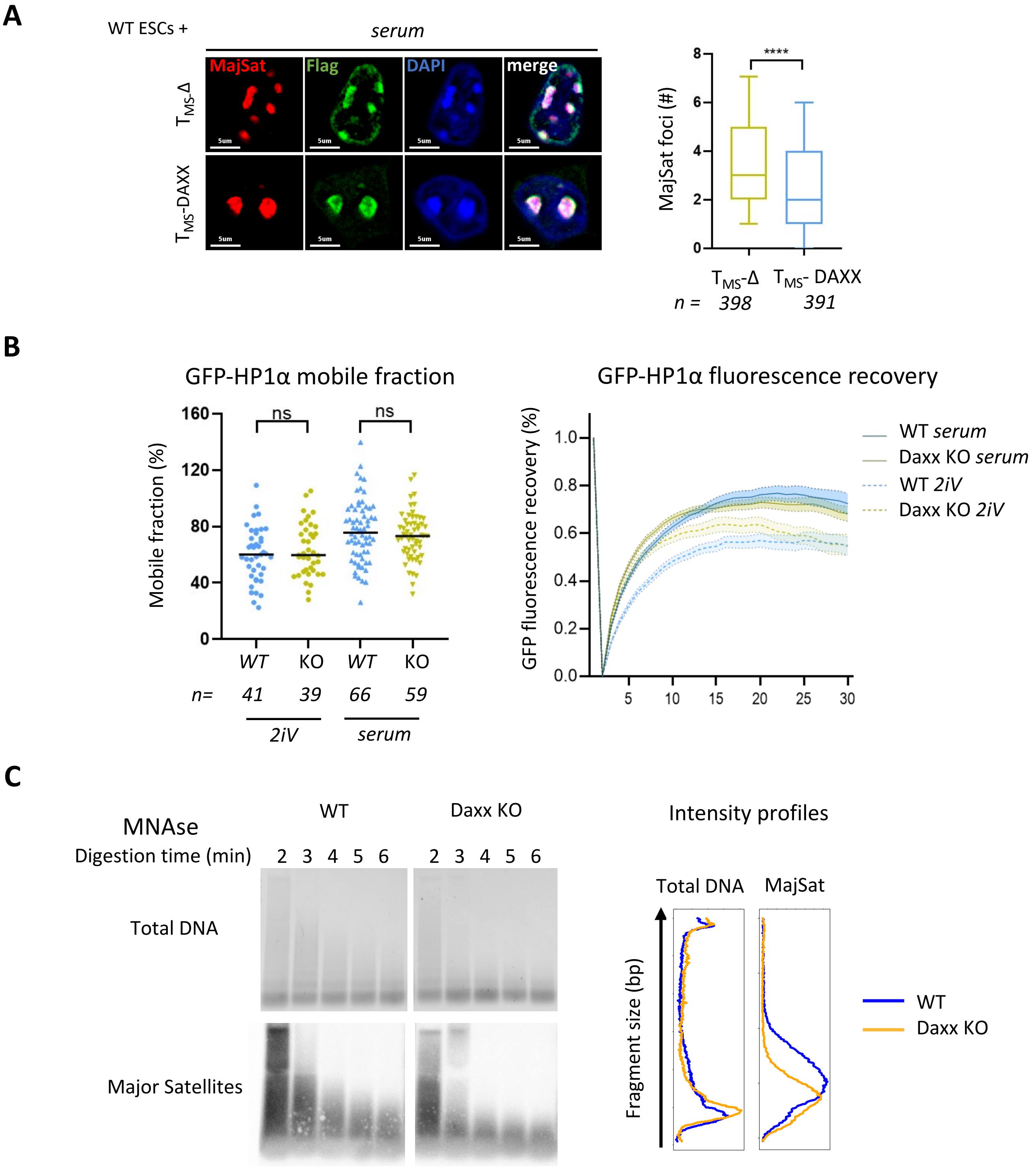


Supplemental Figure S5. Canat et al. 2023

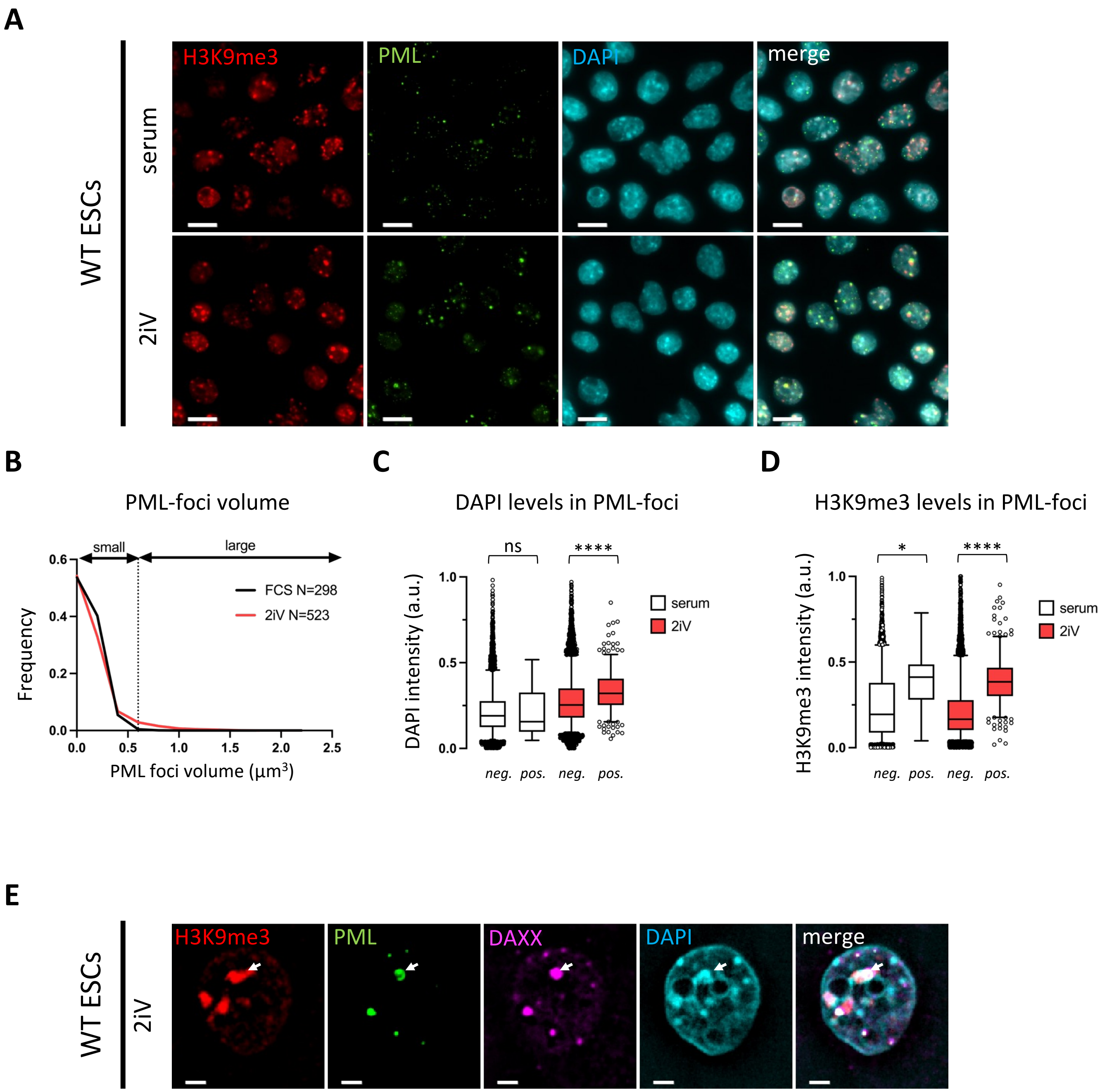


Supplemental Figure S6. Canat et al. 2023

A

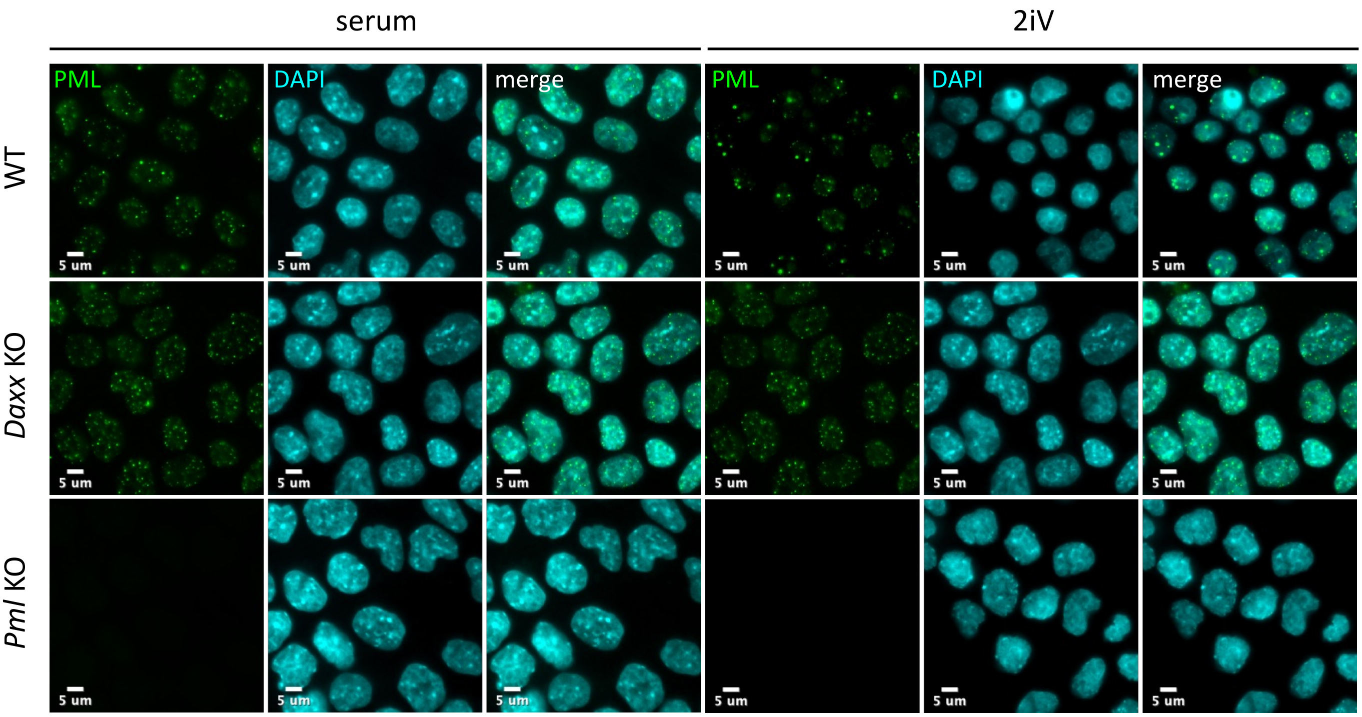


B

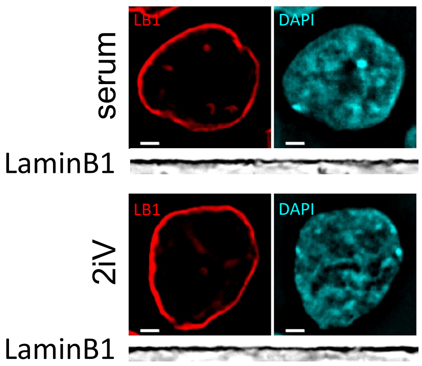


C

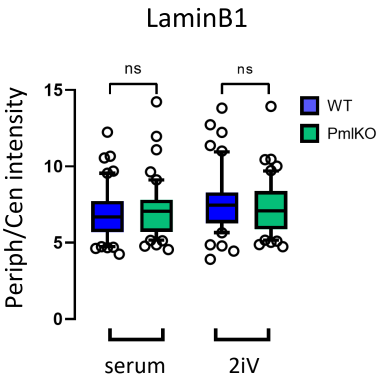


Supplemental Figure S7. Canat et al. 2023

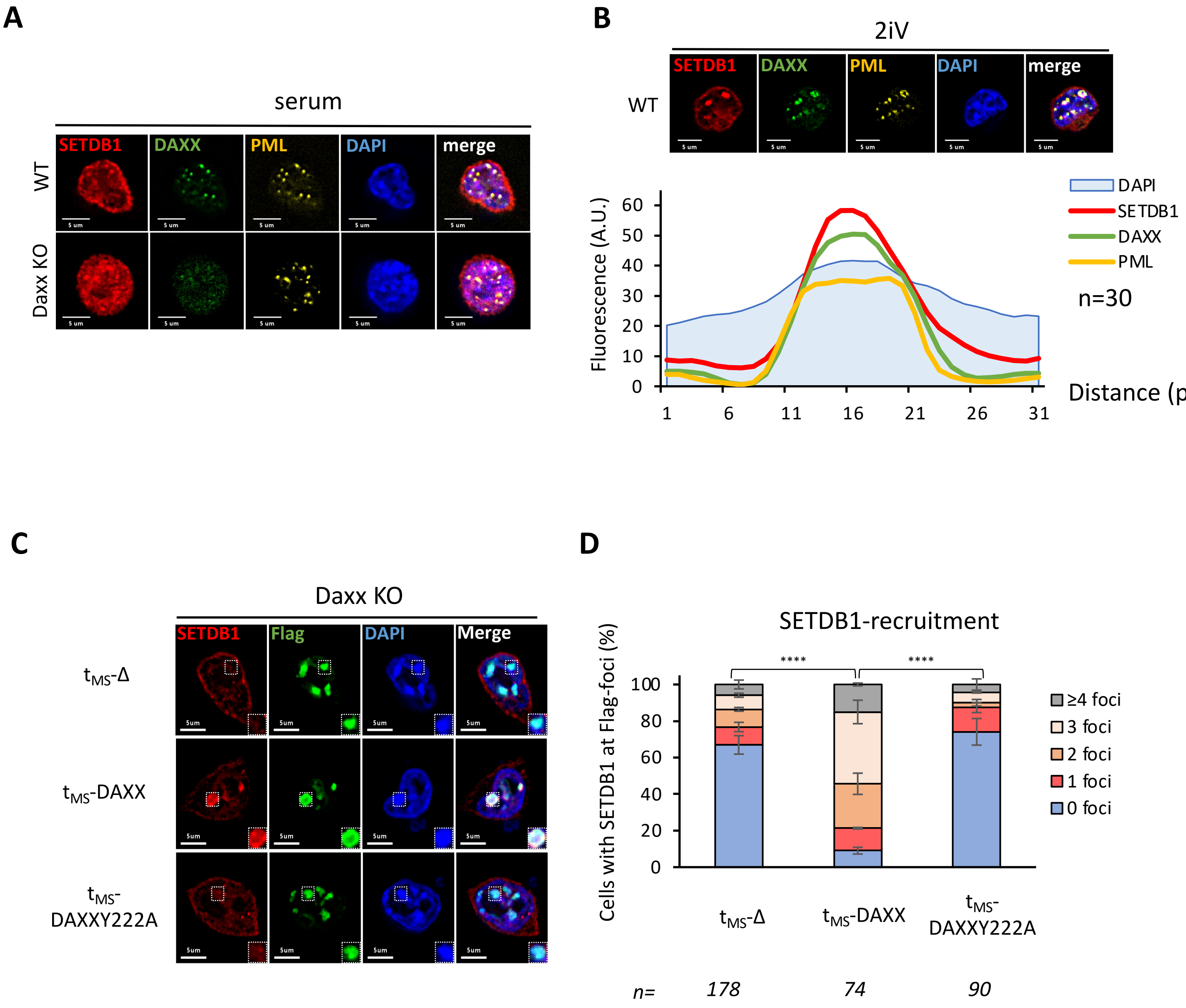


Supplemental Figure S8. Canat et al. 2023

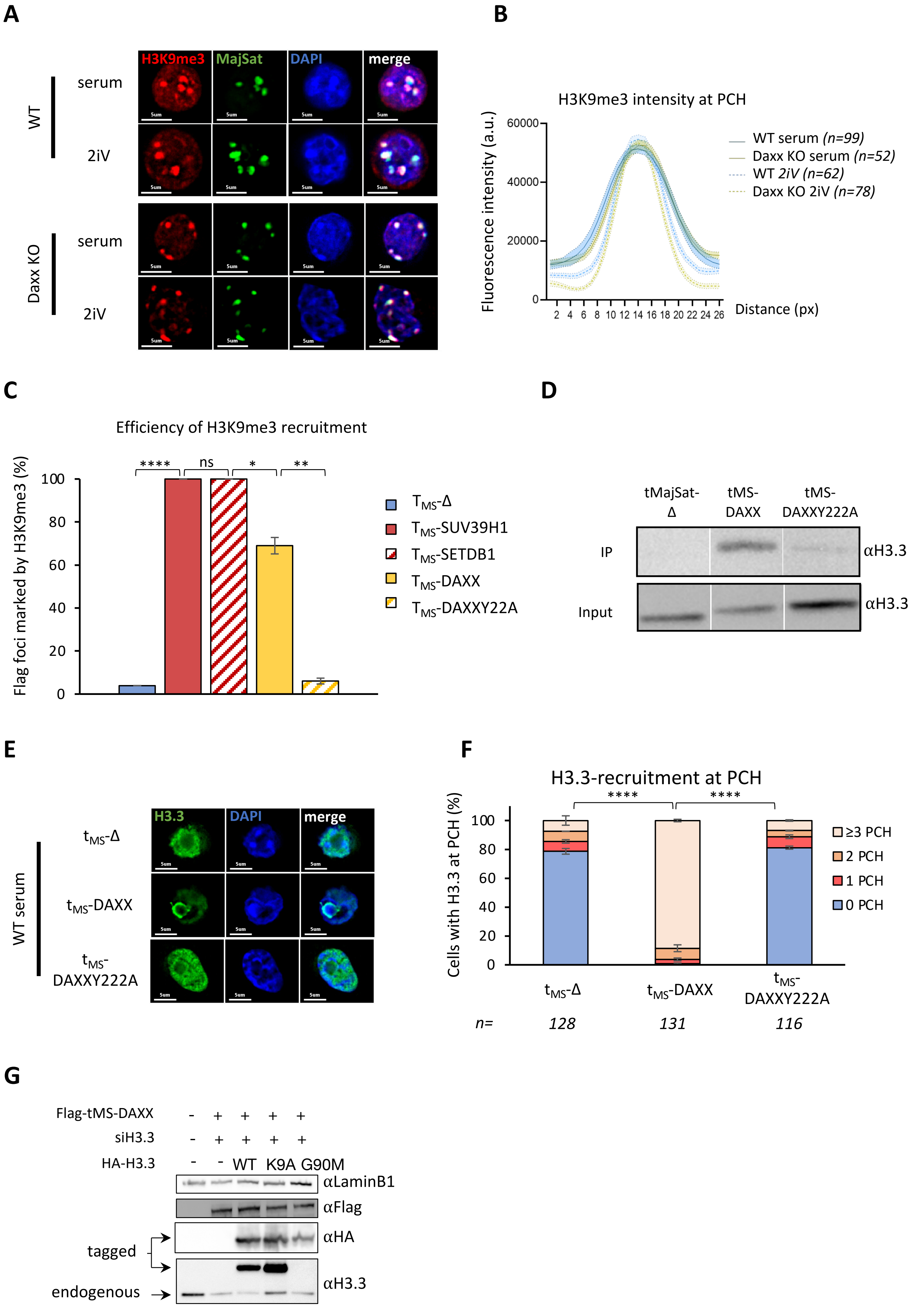


Figure S9. Canat et al. 2023
